# Filopodia rotate and coil by actively generating twist in their actin shaft

**DOI:** 10.1101/2020.09.20.305227

**Authors:** Natascha Leijnse, Younes Farhangi Barooji, Bram Verhagen, Lena Wullkopf, Janine Terra Erler, Szabolcs Semsey, Jesper Nylandsted, Amin Doostmohammadi, Lene Broeng Oddershede, Poul Martin Bendix

**Affiliations:** Niels Bohr Institute, University of Copenhagen, 2100 Copenhagen, Denmark; Biotech Research and Innovation Centre (BRIC), University of Copenhagen, Ole Maaløes Vej 5, 2200 Copenhagen, Denmark; Danish Cancer Society Research Center, Strandboulevarden 49, University of Copenhagen, 2100 Copenhagen, Denmark

## Abstract

Filopodia are actin-rich structures, present on the surface of practically every known eukaryotic cell. These structures play a pivotal role in specific cell-cell and cell-matrix interactions by allowing cells to explore their environment, generate mechanical forces, perform chemical signaling, or convey signals via intercellular tunneling nano-bridges. The dynamics of filopodia appear quite complex as they exhibit a rich behavior of buckling, pulling, length and shape changes. Here, we find that filopodia additionally explore their 3D extracellular space by combining growth and shrinking with axial twisting and buckling of their actin rich core. Importantly, we show the rotational dynamics of the filamentous actin inside filopodia for a range of highly distinct and cognate cell types spanning from earliest development to highly differentiated tissue cells. Non-equilibrium physical modeling of actin and myosin confirm that twist, and hence rotation, is an emergent phenomenon of active filaments confined in a narrow channel which points to a generic mechanism present in all cells. Our measurements confirm that filopodia exert traction forces and form helical buckles in a range of different cell types that can be ascribed to accumulation of sufficient twist. These results lead us to conclude that activity induced twisting of the actin shaft is a general mechanism underlying fundamental functions of filopodia.

## Introduction

Mechanical interactions between cells and their environment are essential for cellular functions like motility, communication, and sensing. The initial contact formed by cells is mediated by F-actin rich filopodia which are highly dynamic tubular structures on the cell surface that allow cells to reach out and interact with their extracellular environment and adjacent cells.^1^

Filopodia are present in wide variety of cell types ranging from embryonic stem cells^2^ to neuronal cells,^3–6^ they are important for cell migration during wound healing^7^ and in cellular disorders such as cancer.^8^ Recently, filopodia have been discovered to play critical roles in development by facilitating communication between mesenchymal stem cells^2^ and during compaction of the early embryo.^9^

Structurally, filopodia are formed as thin membrane protrusions (diameter between 100 and 300 nm) containing 10-30 bundles of actin filaments, which are cross-linked by molecules such as fascin.^10^ Transmembrane integrins link the actin to the cell membrane while peripheral proteins like IBAR link the actin to the inner leaflet of the plasma membrane. ^1^

Filopodia can differ significantly in length from a few micrometers to tens of micrometers. Long specialized filopodia can be sub-categorized into: Cytonemes^11^ which are involved in long range cell signalling; tunneling nanotubes ^2^ involved in intercellular material exchange including cell-cell virus transmission; ^12^ and recently discovered airinemes^13^ found on skin resident macrophages that are during zebrafish development.

Despite the high diversity of mechanical functions carried out by filopodia there seem to be common characteristics which are preserved in all types of filopodia. These include typical traction forces of tens of piconewtons, ^14–18^ growth and shrinkage,^3^ and bending or lever arm activity.^19^

Growth and shrinkage of filopodia are regulated by actin polymerization at the tip^20^ and myosin activity which contributes to retrograde flow.^18,21^ In addition to this, a sweeping motion of the filopodial tip around the cellular base has been reported^3,22^ and even rotational motion has been indicated in HEK293T cells ^17^ and neuronal cells.^6^ However, these studies have not been able to decipher whether the filamentous actin inside the filopodium, in the following denoted as ‘actin shaft’, performs a spinning motion around its own axis or whether the filopodium performs a circular sweeping motion around the anchor point. Complex movements and helical buckling shapes have been reported together with simultaneous force generation,^6,17,22,23^ thus indicating build-up of torsional twist in the actin shaft of HEK293T cell filopodia. The dynamics of filopodial movement has been shown to be unaffected by inhibition of myosin II.^23^ Knock-down of myosin Va and Vb, which are motors walking in a spiral path on actin filaments, was found to reduce the lateral movements of filopodia tips. ^6,24,25^ However, the generality and underlying mechanisms for all these modes of movement has remained enigmatic.

Here we show that twist generation in the actin shaft can explain several of the observed phenomena such as helical buckling, traction generation, and rotational movements of filopodia. We developed a novel optical tweezers/confocal microscopy assay to visualize the rotation of the F-actin shaft inside a filopodium. An optical trap is used to fix the filopodial tip and also used to attach a bead to the side of a filopodium. This bead is indirectly linked to the actin inside the filopodium via vitronectin-integrin bonds and hence reports about any axial rotation and retrograde flow of actin. The observed axial rotations of the bead are compared to results from tracking of filopodial tips of cells grown on glass or embedded in a collagen I matrix. Furthermore, helical buckles resulting from twist accumulation are detected in a number of cell types, on glass as well as embedded in collagen I. To test whether these buckles originate from membrane induced compressional load^26^ or from accumulation of twist within the actin shaft resulting from the spinning, we quantify the forces acting along the actin shaft. The generality of these phenomena was assessed by measuring the traction forces of both naive pluripotent stem cells and terminally differentiated cells. Interestingly, despite the presence of traction forces delivered by filopodia in all cell types, we frequently observed helical buckling in the actin shaft which can be explained by build-up of torsional twist.

We identify a mechanism behind the twisting and consequent spinning of the actin core by modeling actin/myosin complexes inside the filopodial membrane as an active gel, showing that mirror-symmetry breaking and the emergence of twist are generic phenomena in a confined assembly of active filaments, suggesting the physical origin of the observed twisting motion in variety of cell types.

## Results

### Filopodia and extracted membrane tethers can rotate, pull, and buckle

Filopodia show rich and complicated dynamics that is connected to remodeling of their actin shaft. On the order of minutes, they undergo significant movement and reshaping. Filopodial tips are often found to undergo a rotation^6,17,22,27^ or a sweeping motion, see Figure 1A and B for a EGFP Lifeact-7 expressing HEK293T cell on glass and SI movie 1a. Bending or coiling are frequently observed phenomena, shown in Figure 1C and D for a KPC cell (cyan, mEmerald Lifeact-7) embedded in a matrix containing 4 mg/ml collagen I and in SI movie 1b. The traction force generated by filopodia is examined by extracting a new filopodium from the plasma membrane using an optically trapped bead, see Figure 1E-G, and the trapping of the tip also reveals frequent formation of buckles can be observed along the filopodium, see SI movie 1c. Buckle formation also took place without prior imaging with the confocal scanning laser, thus excluding that buckling is an artifact of the cell reacting to the laser illumination, see Figure S1.

**Figure 1:**
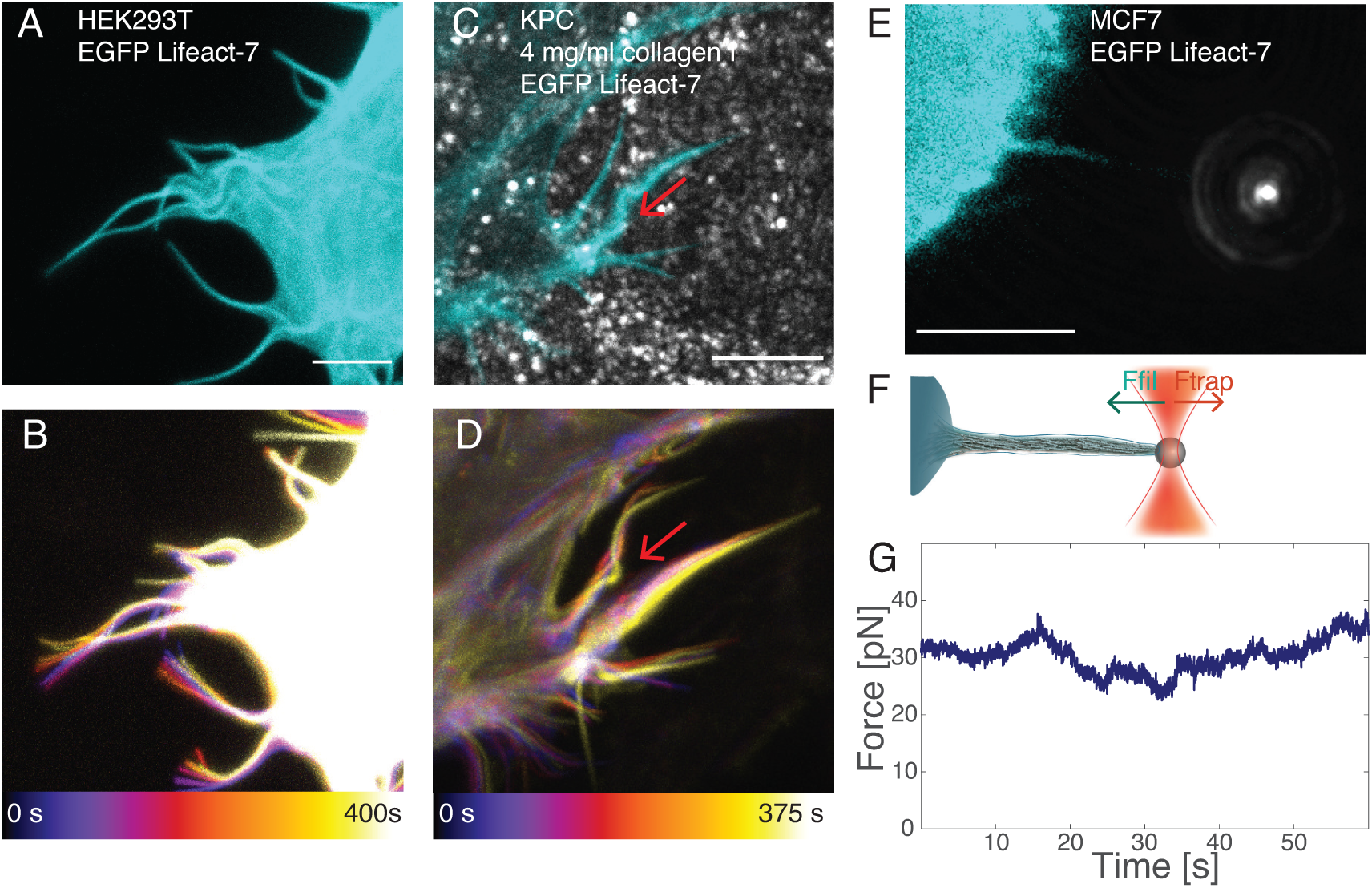
Free and confined filopodia show rich dynamics that can lead to bending, buckling, coiling, shortening, and pulling. (A and B) The tips of free filopodia rotate. Confocal *z*-stacks of a HEK293T cell (cyan, EGFP Lifeact-7) grown on glass. (A) Confocal *z*-projection of a single *z*-stack. Scale bar is 5 *μ*m. (B) Overlay of *z*-projections at 11 consecutive time points (total time is 400 s, color coded from blue to white). (C and D) Filopodia confined by collagen frequently exhibit buckles. Confocal *z*-stacks of a KPC cell (cyan, EGFP Lifeact-7) grown in 4 mg/ml collagen I (grey, reflection). The red arrows mark filopodial bending regions. (C) Confocal *z*-projection of a single *z*-stack. Scale bar is 5 *μ*m. (D) Overlay of *z*-projections at 6 consecutive time points (total time is 375 s, color coded from blue to white. (E, F, G) Membrane tethers extracted from cells fill up with F-actin (actin shaft), are highly dynamic, and behave like filopodia. (E) Confocal image of a tether extracted from a MCF7 cell (cyan, EGFP Lifeact-7) using an optically trapped vitronectin coated *d* = 4.95 m bead (grey, reflection). Scale bar is 5 *μ*m. (F) Schematics of the setup for tether extraction: The force exerted by the optical trap (orange laser beam profile) holds a bead (grey) in place. This bead is used to extract a membrane tether from the cell (cyan). The tether exerts a pulling force on the bead (green arrow) towards the cell body, counteracted by the trap force (red arrow). The force exerted by the tether is measured by tracking the bead’s displacement relative to the initial center of the trap. Over time, the tether begins to bend, coil, and hence shortens and exerts a traction force on the trapped bead. (G) Holding force as a function of time exerted by the tether from (E) versus time.

The rotating motion of a filopodium and the associated shape changes and generation of traction, shown in Figure 1, indicate that the actin shaft contains twist that can be converted into buckling and traction. We therefore next sought to investigate possible generation of torque by an acto-myosin system confined within a tether extracted from a living cell.

### The actin shaft within extracted membrane tethers performs a spinning motion

Filopodia-like structures were formed by extraction of membrane tethers from the cell surface by using optical tweezers.^17–19,28–30^ Depending on the time of observation F-actin can be found to be present inside pulled membrane tethers which subsequently behave like filopodia.^3,4,8,17,31,32^ To visualize and quantify any possible actin shaft rotation we use the setup schematically shown in Figure 2A. A tether, seen as an artificially extended filopodium, was extracted from an EGFP Lifeact-7 expressing cell (as a marker for F-actin^33^) using an optically trapped vitronectin (VN) coated bead (*d* = 4.95 *μ*m). Vitronectin binds to the transmembrane integrins which indirectly attach to the cytoskeletal filamentous actin. Following extraction of the membrane tether the bead was immobilized on the glass surface of the chamber to hold the hereby newly formed filopodium in place. We then used the optical trap to attach another smaller VN coated red labelled bead (*d* = 0.99 *μ*m, in the following denoted as ‘tracer bead’) to the tether and acquired *z*-stacks over time using a confocal microscope. The VN coating on the tracer bead ensures that the bead binds to transmembrane integrin proteins which indirectly interact with actin on the cytosolic side of the membrane. As the actin filaments are known to undergo retrograde flow, so will the integrins and the bead attached to the integrins will move along. Hence, this assay should report the movement of the actin shaft inside the tether.

**Figure 2:**
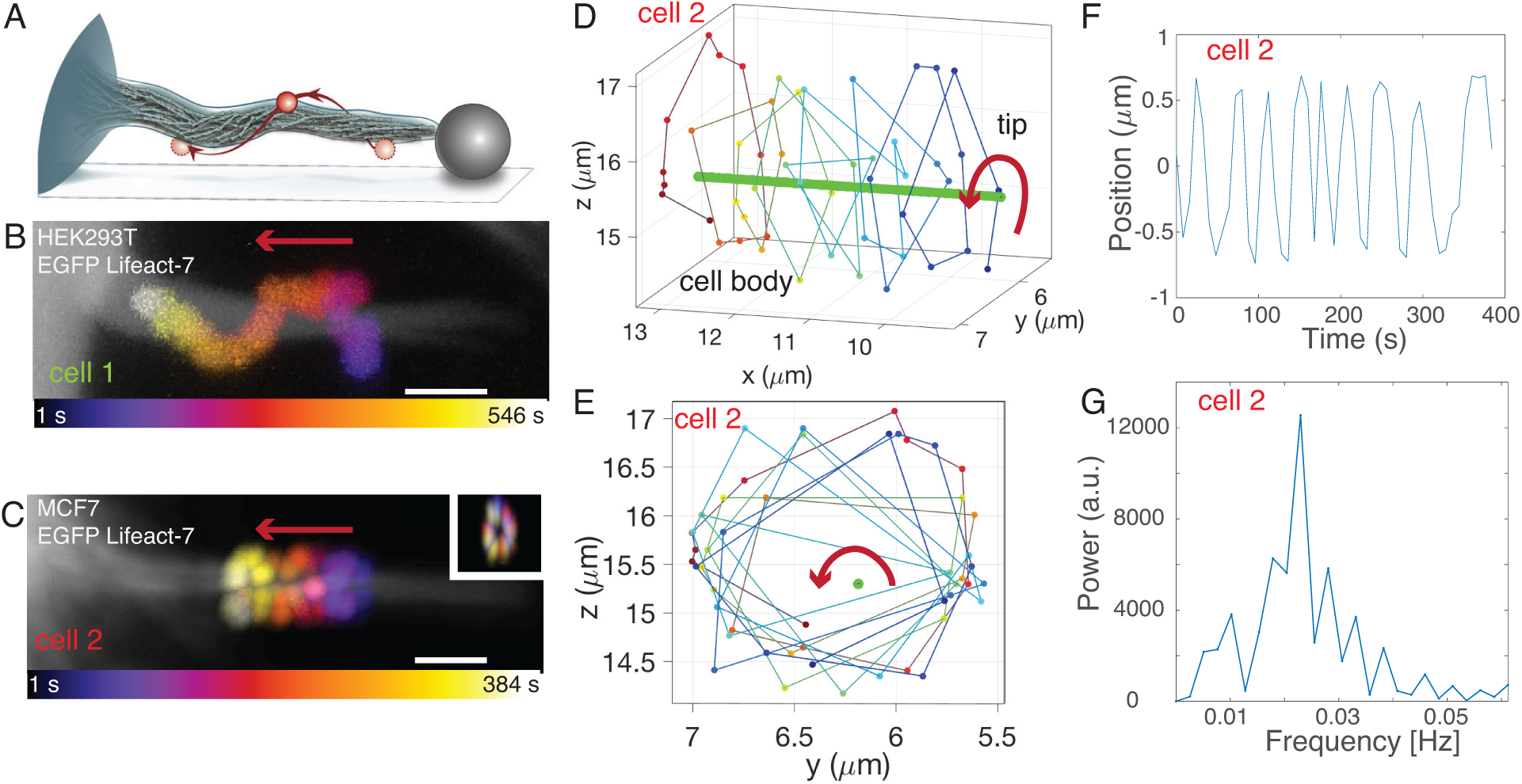
The actin shaft within filopodia performs a spinning motion. (A) Assay to visualize internal F-actin rotation in filopodia: A tether of a cell on a glass slide (gray) is extracted from an EGFP Lifeact-7 expressing cell (cyan) using an optically trapped vitronectin coated bead (grey, *d* = 4.95 m). After tether extraction, the bead is attached to the glass surface of the sample such that the tether is held in place even when the trap is turned off. A tracer bead coated with vitronectin (red, *d* = 0.99 *μ*m) is attached to the tether using the optical trap. The tracer bead binds indirectly to the filamentous actin inside the tether via transmembrane integrins. After the tracer bead is attached, the trap is turned off and confocal *z*-stacks are acquired at consecutive time points revealing the spinning of the F-actin shaft inside the filopodium. (B and C) Confocal *z*-projections of a tether from a HEK293T cell (B, cell 1, grey, EGFP Lifeact-7) and a not activated MCF7-p95ErbB2 cell (C, cell 2, grey, EGFP Lifeact-7) at consecutive time points where an attached tracer bead rotates counterclockwise around the actin shaft from tip towards the cell body (color coded from yellow to red). Scale bars are 2 *μ*m. Red arrows show direction of motion from tip towards cell body. The inset in C shows images of the tracer bead position at all time points in *yz*-view. (D and E) 3D trajectory of the tracer bead (from blue to yellow, same time scale as in C) moving counterclockwise around the filopodium (green) towards the cell body in *xyz* (D) and *yz* (E) view. (F) Tracer bead position as function of time for cell 2. (G) The rotation frequency of the tracer bead and thus the actin shaft rotation frequency is found to be 0.022 Hz, as obtained from the peak of the power spectrum.

Results of this assay are shown in Figure 2B and Figure S2A for a bead attached to a tether from a HEK293T cell (‘cell 1’). Figure 2C-G and SI movie 2a show bead rotation data for ‘cell 2’, an uninduced MCF7-p95ErbB2 breast carcinoma cell expressing a N-terminally truncated 95 kDa version of the p95ErbB2 receptor. ^34,35^

As shown in Figure 2B-D and Figure S2A, the tracer bead moves towards the cell body of the HEK293T (Figure 2B and Figure S2A) and the uninduced MCF7-p95ErbB2 cell (Figure 2C, SI movie 2a) as expected due to the retrograde flow. The transport of the tracer bead along the filopodium was at the speed of 150 ± 131nm/s, see Table S2C and is consistent with retrograde flow of actin within the filopodium. This behavior was also observed for MCF7 cells not expressing the p95ErbB2 receptor, see Figure S2B.

We used a custom written MATLAB program to subtract the constant sideway movement of the filopodium and thereby align the axis of the filopodium in the *yz*plane (see Figure S6). This revealed that the bead furthermore undergoes a spiralling motion around the filopodium. We isolated the rotation of the bead around the axis of the actin shaft, see Figure 2D,E, and found it to be clounterclockwise as seen from tip towards the cell body. The rotation frequency of the VN coated tracer bead attached to ‘cell 2’ in Figure 2C was 0.002 Hz as obtained from a power spectral analysis, see Figure 2F,G. Table S2C shows additional tracer bead rotation frequencies measured for beads on filopodia from HEK293T, MCF7, and uninduced MCF7-p95ErbB2 cells.

The measured spinning of the actin core is also expected to result in rotational motion of the filopodial tip. Therefore, we next tracked the three dimensional movement of free filopodia and quantified their rotary frequencies.

### The tips of filopodia rotate with a similar frequency as the spinning of the actin shaft

We tracked the motion of F-actin labelled filopodia (via EGFP Lifeact-7 expression) in cells grown on glass slides, Figure 3A, and cells embedded in collagen I gels, Figure 3E. The tracking of filopodia was performed using a custom written MATLAB algorithm which allows tracking free tips of filopodia during their growth and shrinkage. The algorithm allowed us to compensate for filopodial drift caused by the migration of the cell and by possible lateral sliding of the filopodium.

**Figure 3:**
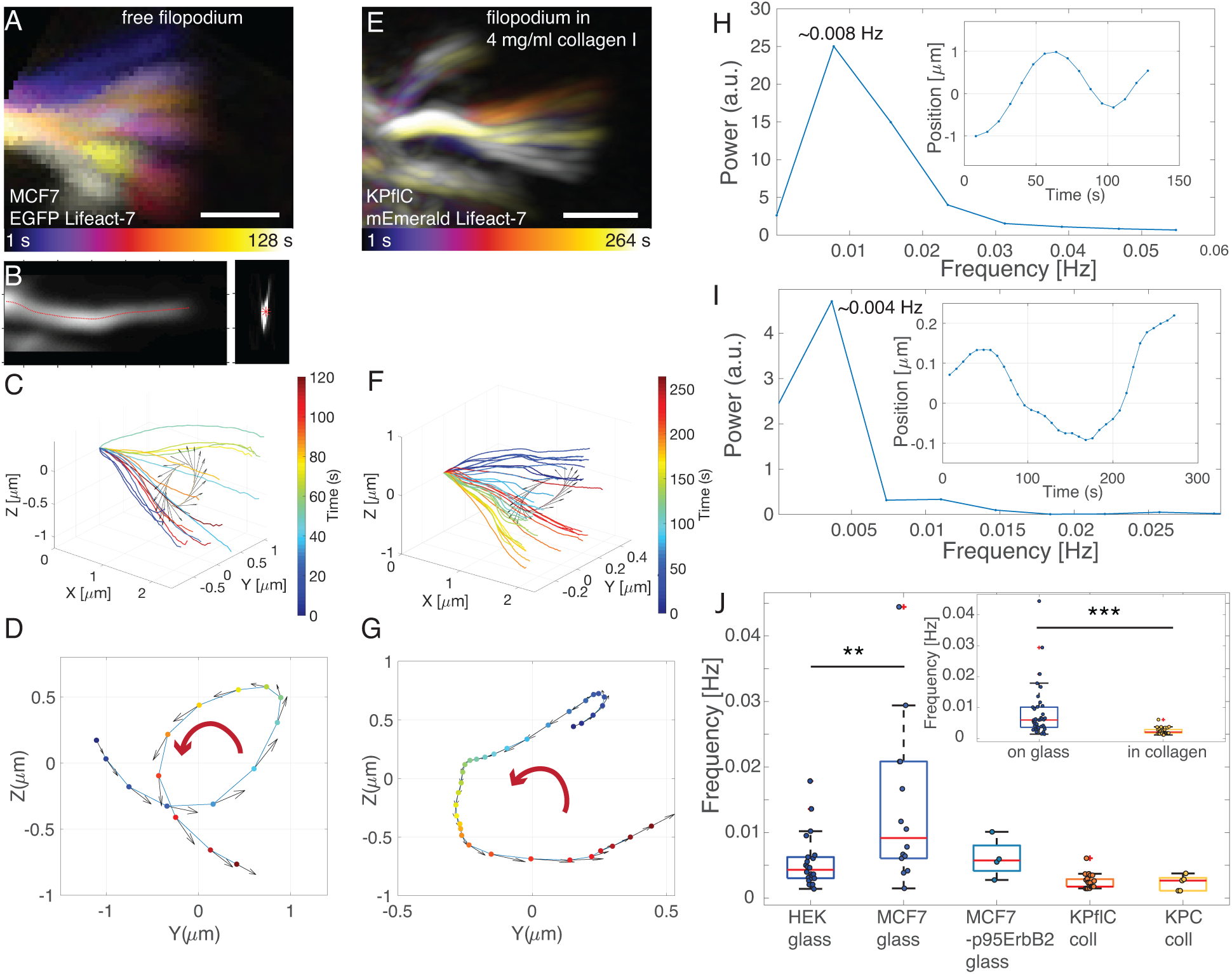
The tips of free filopodia of cells grown on glass and filopodia of cells grown in a collagen I matrix undergo a rotary motion with different frequencies. (A) Gabor-filtered *z*-projection of the filopodial tip movement of a EGFP Lifeact-7 labelled MCF7 cell at 16 consecutive time points for a total time of 128 s. Scale bar is 1 *μ*m. (B) *xz* (left) and *yz* (right) view of a single *z*-projection showing the Gabor filtered data (grey) overlayed with the traced filopodium (red). (C and D) Result from tip tracking of the filopodium from (A) in *xyz* (C) and *yz* view (D). Tip tracking over time reveals a counterclockwise rotation of the whole filopodial tip, seen from tip towards the cell body, as indicated by the red arrow. (E) Gabor-filtered *z*-projection of filopodial tip movement of an EGFP Lifeact-7 labelled KP^fl^C cell grown in 4 mg/ml collagen I at 16 consecutive time points for a total time of 264 s. Scale bar is 1 *μ*m. (F and G) Result from tip tracking of the filopodium shown in (E) in *xyz* (F) and *yz* view (G). Tip tracking over time reveals a counterclockwise rotation of the whole filopodial tip, seen from tip towards the cell body, as indicated by the red arrow. (H and I) The rotation frequency of the filopodial tip, found via power spectral analysis, for the MCF7 cell from (A) is 0.008 Hz (H) and the KP^fl^C cell in 4 mg/ml collagen I from (E) is 0.004 Hz (I). Insets: Filopodial tip position as function of time. (J) The tip rotation frequencies for HEK293T, MCF7 and MCF7-p95ErbB2 cells grown on glass and KP^fl^C and KPC cells grown in 4 mg/ml collagen I differ significantly from each other. Red lines are the medians. Inset: Plot of all tip rotation frequencies on glass (mean ± std: 0.0084 ± 0.0084 Hz) and in collagen I (mean ± std: 0.0024 ± 0.0011 Hz). Mean rotation frequencies and *p*-values can be found in Tables S3B,C.

We acquired confocal *z*-stacks of cells grown on glass and in 4 mg/ml collagen I gels for approximately 5 min. Figure 3A and E show an overlay of *z*-projections of a filopodium from a MCF7 cell cultured on glass and a KP^fl^C cell in 4 mg/ml collagen I, respectively, at different time points. The lateral movement in Figure 3A and E has been subtracted such that filopodia at all times initiated at the same point, and thus their rotary motion could be tracked in 3 dimensions, see Gabor filtered image of the filopodium in Figure 3B shown in *xz* (left) and *yz* view (right).

Since the length of a single filopodium varies over time, we chose a common *yz* plane close to the tip through which the filopodium crossed at all time points. Figures 3C, D and Figure 3F, G show that the filopodial tips rotate counterclockwise over time in the *xy* plane as seen from the tip towards the cell body for the MCF7 cell on glass and the KP^fl^C cell in a collagen I matrix, respectively. We obtained the rotation frequencies via a power spectral analysis, see Figure 3H and I. The tips of filopodia on the MCF7 and KP^fl^C cells rotated with a frequency of 0.008 Hz and 0.004 Hz, respectively. Figure 3J compares median rotation frequencies for HEK293T, MCF7, and uninduced MCF7-p95ErbB2 cells on glass and KP^fl^C and KPC cells in collagen I matrices. Rotation frequencies of filopodial tips in 3D collagen I matrices are clearly lower than of free cells (Figure 3J, inset) indicating that the extracellular fibers confine and slow down the tip movement. The mean values and standard deviations as well as the *p*-values can be found in Tables S3B,S3C.

Of the 74 filopodial tips we found 46 % to undergo a counterclockwise rotation (as seen from tip towards the cell body), 7 % were clockwise. In 47 % of the cases we could not clearly determine a handedness which is due to the fact that filopodial dynamics has many more contributors than actin shaft rotation, such as retrograde flow and overall cell movement. See example trajectories in Figure S3A.

We also detected filopodia rotation in naive pluripotent mouse embryonic stem cells (mESCs) grown in 2i medium, see SI movie 3a. These cells represent the earliest development and hence could indicate whether filopodia rotation is a general property of cells. Comparing these with terminally differentiated cells like Hepa 1-6 mouse hepatocytes (SI movies 3b, c), and invasive cancer cells such as MCF7-p95ErbB2, see Tables S3B,S3C, shows that filopodia rotation takes place in both, the earliest and late stages of development.

We hypothesize that the observed rotary movement of filopodial tips, quantified in Figure 3, is generally caused by a spinning axial rotation of the actin shaft as measured in Figure 2. A clear indicator of axial rotation of actin in filopodia is the presence of helical buckling which arises from over-twisting of the actin shaft. In the following, we therefore focus on filopodia buckling in cells cultured in 3D collagen I networks in which dynamic filopodia can build up twist by interacting with the collagen I fibers.

### Helical buckling and coiling of filopodia

Filopodia from cells grown in 3D collagen I gels dynamically explore their 3D environment, but they are found to exhibit a more restricted motion compared to cells grown on glass due to the confinement of the fibers surrounding the cell, as shown in Figure 4. Figure 4A-E shows a KP^fl^C cell labelled with mEmerald-Lifeact-7, embedded in a 4 mg/ml collagen I gel, imaged with a confocal microscope. Bending, buckling, and helical coiling of the filopodium become apparent in a color-coded confocal *z*-stack at 3 consecutive time points over a time span of 72 s, see also SI movie 4a.

**Figure 4:**
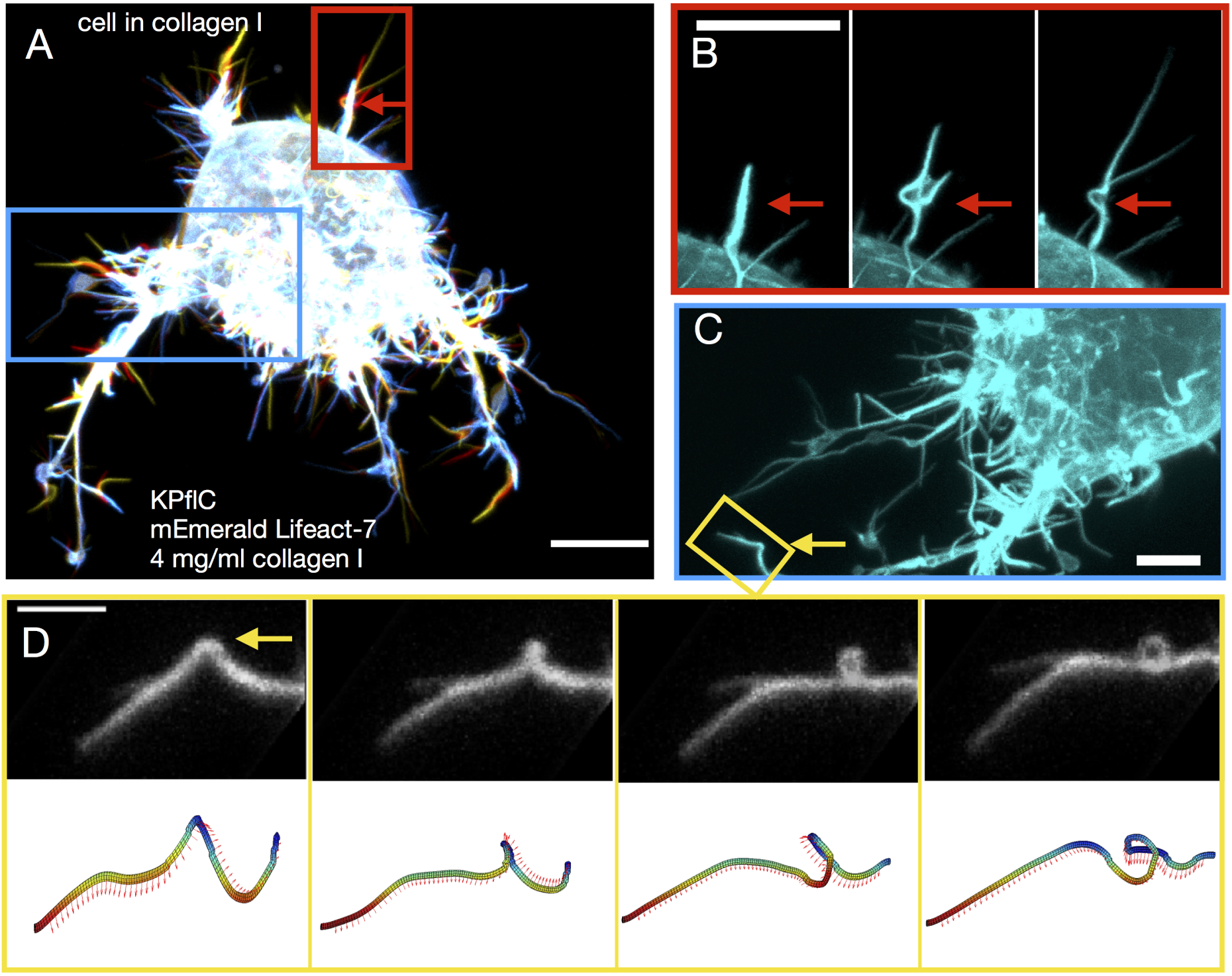
Filopodia of cells embedded in a collagen I matrix undergo helical buckling as a result of rotation and twist accumulation upon contact with the matrix. (A) Overlay of 3 confocal *z*-projections of a KP^fl^C cell (mEmerald Lifeact-7) embedded in 4 mg/ml collagen I at consecutive time points (color coded over 72 s). The red arrow highlights a region where a buckle forms. Scale bar is 10 *μ*m. (B) Zoom-in on the red rectangular region from (A): Individual images of the 3 time points show buckle formation. Scale bar is 10 *μ*m, frame interval is 24 s. (C) Zoom-in on the blue rectangular region from (A) (rotated). The yellow rectangle highlights a bending filopodium. Scale bar is 5 *μ*m. (D) Top: Gabor-filtered images of a the filopodium from the yellow region in (C) at 4 consecutive time points (frame interval is 19 s) of a KP^fl^C cell in 4 mg/ml collagen I. Scale bar is 2.5 *μ*m. Bottom: 3D tracks of the filopodium show bending and coiling. Red arrows indicate the binormal vectors along the filopodium.

Filopodia growing in a 3D collagen I network will experience external friction at the contact points between filopodium and the fibers. Additionally, friction exists between the plasma membrane and the actin shaft mediated by various proteins linking the membrane with the actin. The helical buckles and coils observed in Figures 4B, C and D, S4A and SI movie 4b can hence be a signature of over-twisting of the actin shaft which occurs naturally when rotating filopodia interact with a collagen I network or the membrane.

### Filopodia buckle and pull at the same time

To further investigate the nature of these buckles we tested whether such a twist-buckling transition occurs in presence of traction in the filopodium. We therefore measured the traction force when holding the membrane tether by an optical trap, as shown schematically in Figure 5A. Compressive buckling from membrane tension, as suggested in references^26,36^ is an unlikely mechanism if the filopodium is able to exert a traction force while undergoing buckling. We observed buckling in filopodia, without the presence of external collagen I fibers, in optically trapped filopodia pulled from cells cultured on glass, see Figure S4B and SI movie 4b.

**Figure 5:**
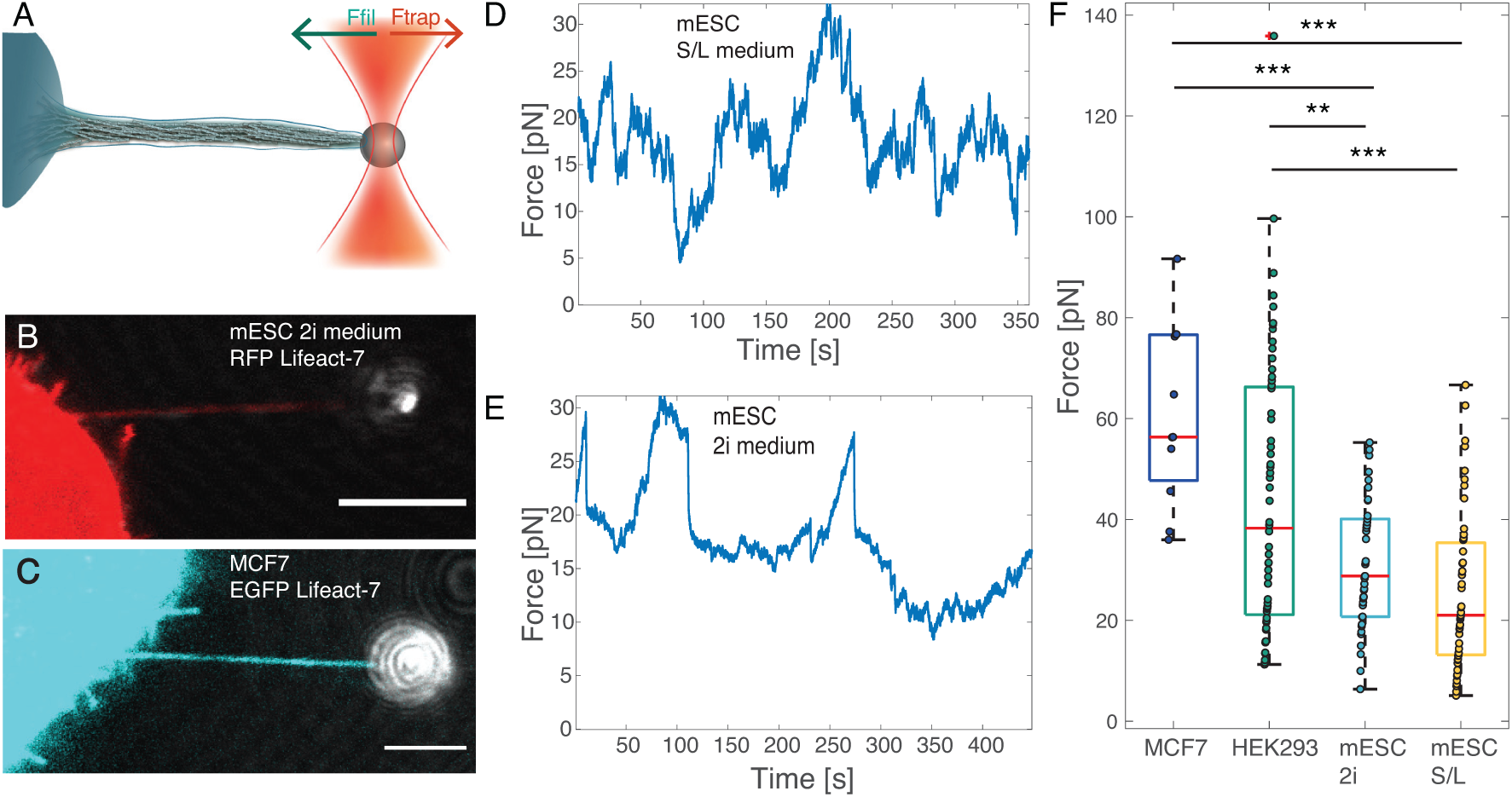
Tethers pulled from different cell types– from naive pluripotent stem cells to terminally differentiated cells– are highly dynamic and generate significant traction forces. (A) Schematics of the optical tweezers assay to measure the dynamic force exerted by a tether. An optically trapped VN coated bead (*d* = 4.95 *μ*m) is used to extract and hold a membrane tether at a given length with a holding force *F*_*trap*_. (B) Confocal image of a filopodium extracted from a mESC (red, RFP Lifeact-7) grown in 2i medium. Scale bar is 5 *μ*m. (C) Confocal image of a filopodium extracted from a MCF7 cell (cyan, EGFP Lifeact-7). Scale bar is 5 *μ*m. (D) and (E) Force curves for tethers pulled from mESCs grown in 2i (D) and in S/L (E) medium. (F) Maximum tether holding forces measured for mESCs cultured in 2i and S/L medium plotted together with maximum forces measured for MCF7 and HEK293 cells. From left to right: MCF7 (N = 11, mean ± std = 61.10±17.85), HEK293 (N = 60, 43.88 ± 27.14), mESC in 2i medium (N = 42, 31.32 ± 12.73), and mESC cells in S/L medium (N = 51, 25.56 ± 15.40). Red lines mark the medians. (* p≤ 0.05, ** p≤ 0.01, *** p≤ 0.001). Mean, maximum forces, and *p*-values are shown in Tables S5C,D.

We extracted tethers from mouse embryonic stem cells and terminally differentiated cells (MCF7) that were transfected to express EGFP Lifeact-7 labelled F-actin, by optical trapping of a vitronectin coated bead, see Figure 5B, C, respectively. After extracting a membrane tether with a trapped bead, small amounts of F-actin were immediately present inside the tether, but over a time course of ∼ 150 s, the cell recruited more F-actin, as seen in Figure S5A for a mESC, and the actin shaft was observed to extend several micrometers into the membrane tube. Subsequently, the new structure became highly dynamic and behaved like a filopodium, see also SI movies 5a,b. The force exerted by the filopodium in Figure S5A was measured over a time span of 60 s by tracking the bead’s displacement from the initial center of the trap.

To investigate the general traction force delivered by cells existing in different developmental stages we quantified the maximal filopodium traction forces exerted by MCF7, HEK293 cells, and mESCs cultured in 2i or Serum/LIF (s/L) medium, see Figure 5D-F. We found that all cell types exclusively exert a traction force in the range 20-80 pN on the trapped bead and stem cells cultured in 2i or S/L medium were found to exert slightly lower maximal forces. Mean forces and standard deviations and *p*-values can be found in Tables S5C,D. Time traces of forces from MCF7 cells in Figure S5B show that the extracted tethers display similar activity as expected for filopodia. The high traction forces exceed the typical force of 10 pN needed for holding a pure membrane tether containing no F-actin.^4,18^ Since buckles can be observed in force generating tethers held by an optical trap we conclude that compressive forces from the membrane are unlikely to be responsible for the observed buckling and coiling of filopodia, but instead our data suggest that the actin structure exhibits an internal twist generating mechanism.

### Twist deformations are a generic non-equilibrium feature of confined actomyosin complexes

Our experimental observations demonstrate that the complex dynamics of filopodia including helical rotation, buckling, and tip movement are induced by the twisting motion of actin filaments inside the filopodia. Furthermore, the emergence of such twisting motion in a variety of cell types and in both early and late stages of development points to a possible generic mechanism for the formation of twist in actin/myosin complexes within the filopodial cell membrane. In order to understand the underlying mechanism of twist generation, we next used a three-dimensional active gel model to study the dynamics of actin/myosin complexes confined within a geometrical constraint. This choice was motivated by the generality of this class of continuum models which is due to the fact that only local conservation laws, interaction between the systems’ constituents, and perpetual injection of energy at the smallest length-scale are assumed. Furthermore, active gel equations have proven successful in describing several aspects of the physics of actin/myosin networks including actomyosin dynamics at the cell cortex,^37,38^ actomyosin induced cell motility,^39,40^ actin retrograde flows,^41,42^ and topological characteristics of actin filaments.^43,44^ Within this framework the dynamics of actin/myosin complexes are expressed through a continuum representation of their minimal degrees of freedom including the orientation and velocity fields. The orientation field is represented by a tensor order parameter **Q** = 3*q/*2 × (*nn*^T^ − **I***/*3), where *q* is the magnitude of the orientational order and *n* is the director, representing the coarse-grained orientation of the actin filaments.^45,46^ The dynamics of the orientation tensor **Q** follows Beris-Edwards equations,^47^ describing the alignment to flow and relaxation dynamics due to the filament elasticity *K*. The orientation dynamics is coupled to the velocity field governed by a generalized Stokes equation that accounts for the active stress generation due to force dipoles associated with actin tread-milling, described as **Π**^active^ = −*ζ***Q**, where *ζ* denotes the strengths of the active stress generation (see *Supplementary Information* for the details of the governing equations and the mapping of parameters to physical units).

In order to emulate the dynamics of actin filaments inside the filopodia, we consider a simplified setup of an active gel confined inside a three-dimensional channel with a square cross-section of size *h*. In two-dimensions it is shown that increasing the confinement size above a threshold results in a spontaneous shear flow inside the confinement due to bend-splay deformations. ^48,49^ Interestingly, we find that here increasing the size of the three-dimensional confinement beyond a threshold results in the spontaneous flow generation accompanied by the emergence of twist deformations (Figure 6A,B). To characterize the amount of the twist in the system we measure the average twist deformations across the channel 𝒯 = ⟨|*n* · (∇ × *n*) |⟩_*x,t*_, where ⟨ ⟩_*x,t*_ denotes averaging over both space and time. As evident from Figure 6C increasing the confinement size above a threshold results in increasing amount of twist in the active gel. The amount of the twist generation and the threshold are further controlled by the activity of the filaments *ζ* and their orientational elasticity *K*. For a fixed channel size *h* increasing activity triggers a hydrodynamic instability^48,50^ that results in spontaneous flow generation and twist deformations (Figure 6D) consistent with the experimental observations in Figure 4 and Figure S8. The hydrodynamic instability and the creation of the spontaneous flows are suppressed by the filament elasticity *K*. As such increasing the filament elasticity reduces the amount of twist in the system up to a threshold value, where any twist generation is completely suppressed (Figure 6E).

**Figure 6:**
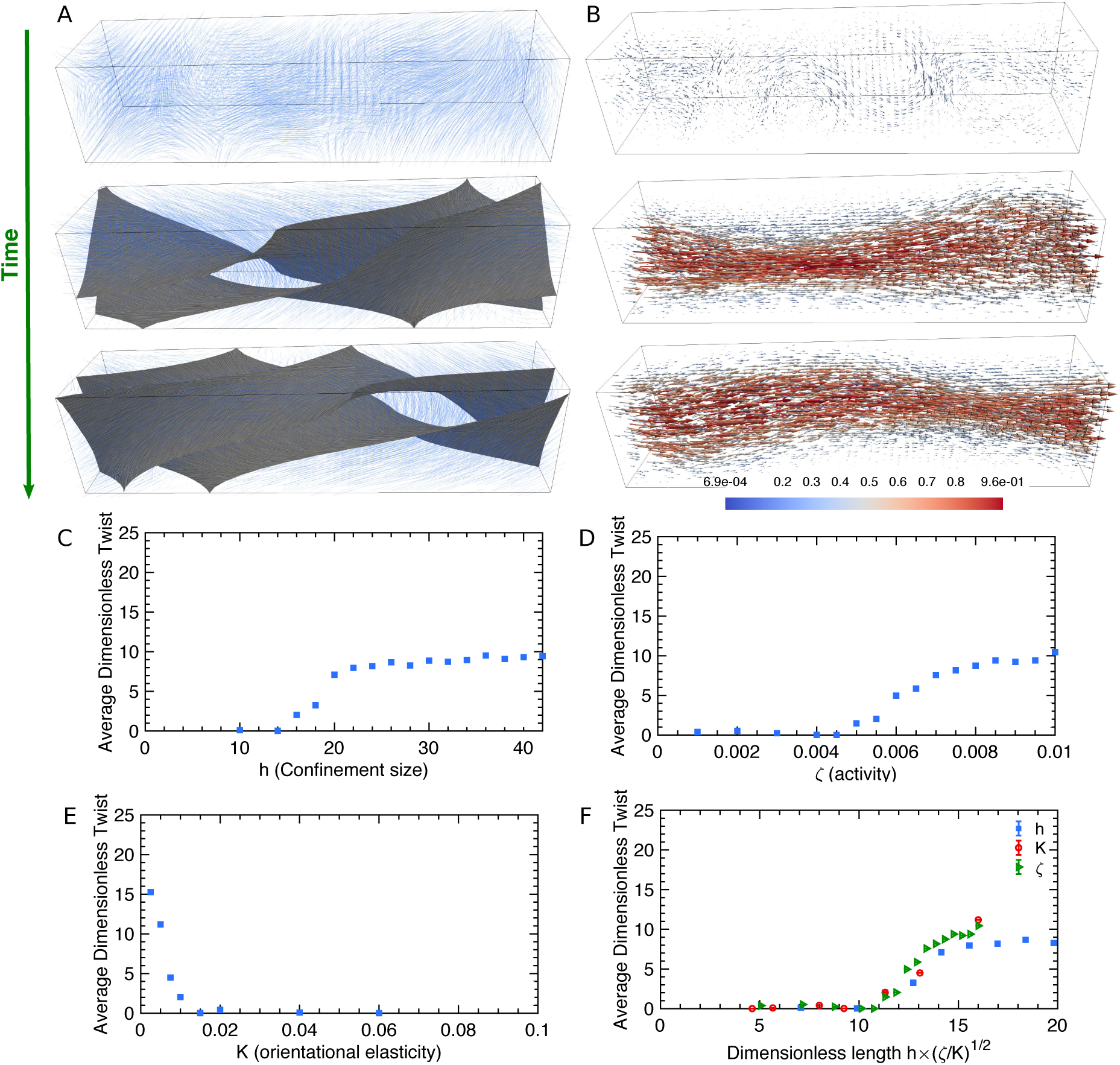
Actin shaft twisting is a generic phenomenon caused by the activity of actomyosin complexes inside the confining filopodial cell membrane. (A,B) Temporal evolution of (A) the orientation and (B) velocity fields of the active gel representing actin filaments/myosin motor mixtures. In (A) the director field associated with the coarse-grained orientation of actin filaments 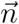 is shown by blue solid lines, and is overlaid with the isocontours of twist deformations 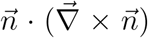. In (B) the colormap indicates the magnitude of the velocities normalized by the maximum velocity. (C,D,E) Dependence of the average twist on (C) the confinement size, (D) activity, and (E) elasticity. The average amount of twist is non-dimensionalized by the channel length. (F) Average twist as a function of the dimensionless length, for varying confinement sizes, activities, and elasticities, showing the collapse of the data.

The interplay between the activity, elasticity, and the confinement size can be best understood in terms of a dimensionless parameter 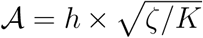 which describes the competing effect of two length scales: the confinement size *h* and the length scale set by combined effects of activity and elasticity 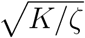. Indeed plotting the amount of twist as a function of the dimensionless number 𝒜 results in the collapse of the data corresponding to varying activity, elasticity, and confinement size (Figure 6F), indicating that 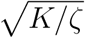 is the relevant activity-induced length scale: increasing the activity enhances active stress generation and orientational deformations, while such deformations are accommodated by the elasticity. As a result, larger activity (or smaller elasticity) leads to the emergence of deformations with smaller length scales. On the other hand, suppression of the activity is expected to increase the deformation length scale. When this length scale becomes larger than the confinement size, all deformations are suppressed and no twist is expected in the actin filaments.

### Chirality of twist

It is important to note that the emergence of twist happens without having any explicit chirality in the equations of motion and is due to a hydrodynamic instability that is induced by the activity of the confined actomyosin complexes. Since this hydrodynamic instability breaks the mirror-symmetry, it will choose both clockwise or counterclockwise directions of rotation with a similar probability. From a biological perspective there could be several contributors for biasing the orientation of the twist. The molecular motors myosin V^24^ and myosin X^51^ have been shown to walk along a spiral path on actin bundles found in filopodia. These motors walk toward the tip in a counterclockwise orientation which would contribute to a torque in the opposite direction. Our experiments suggest that the rotation of filopodia is biased towards the opposite orientation of the path formed by these myosin motors which could suggest that motors increase the probability for the measured orientation. To capture this experimentally measured bias in our active gel dynamics we refine our model to include active stress originating from torque dipoles.

The effect of torque dipoles is accounted for through additional contributions to the active stress 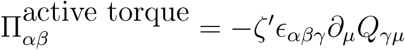, where *ϵ*_*ijk*_ is the Levi-Chevita operator and *ζ*′ controls the strength of the torque dipole.^37,52^ It is easy to see that this term already breaks the mirror-symmetry, such that positive (negative) *ζ*′ exerts clockwise (counterclockwise) torque. No experimental measure of the relative strength of force and torque dipoles are available, however, dimensional analyses suggests that the ratio of active stress coefficients *ζ*′*/ζ* ∼ 0.1 indicating that in general for actomyosin complexes the contributions of torque dipoles to the dynamics are small compared to the force dipoles (see Supplementary Information for the estimate of the torque dipole coefficient). Furthermore, the emergence of small, but finite number of clockwise rotations in the experiments, show that the hydrodynamic instability is the controlling mechanism for the mirror symmetry breaking. Nevertheless, even small values of this chiral terms will bias the mirror-symmetry breaking in the direction of the torque dipole. Indeed our representative simulations show how changing the sign of *ζ*′ results in the change in the handedness of the filaments rotation around the axis of the channel (see Figure S9).

## Discussion

Our results show that axial twisting and filopodia rotation is generic in cells and is detected in naive pluripotent stem cells and in terminally differentiated cells. The spinning of the actin shaft leads to a rich variety of physical phenomena such as tip movement, helical buckling, and traction force generation and this allows the cell to explore the 3D extracellular environment while still being able to exert a pulling force as summarized in Figure 7.

**Figure 7.**
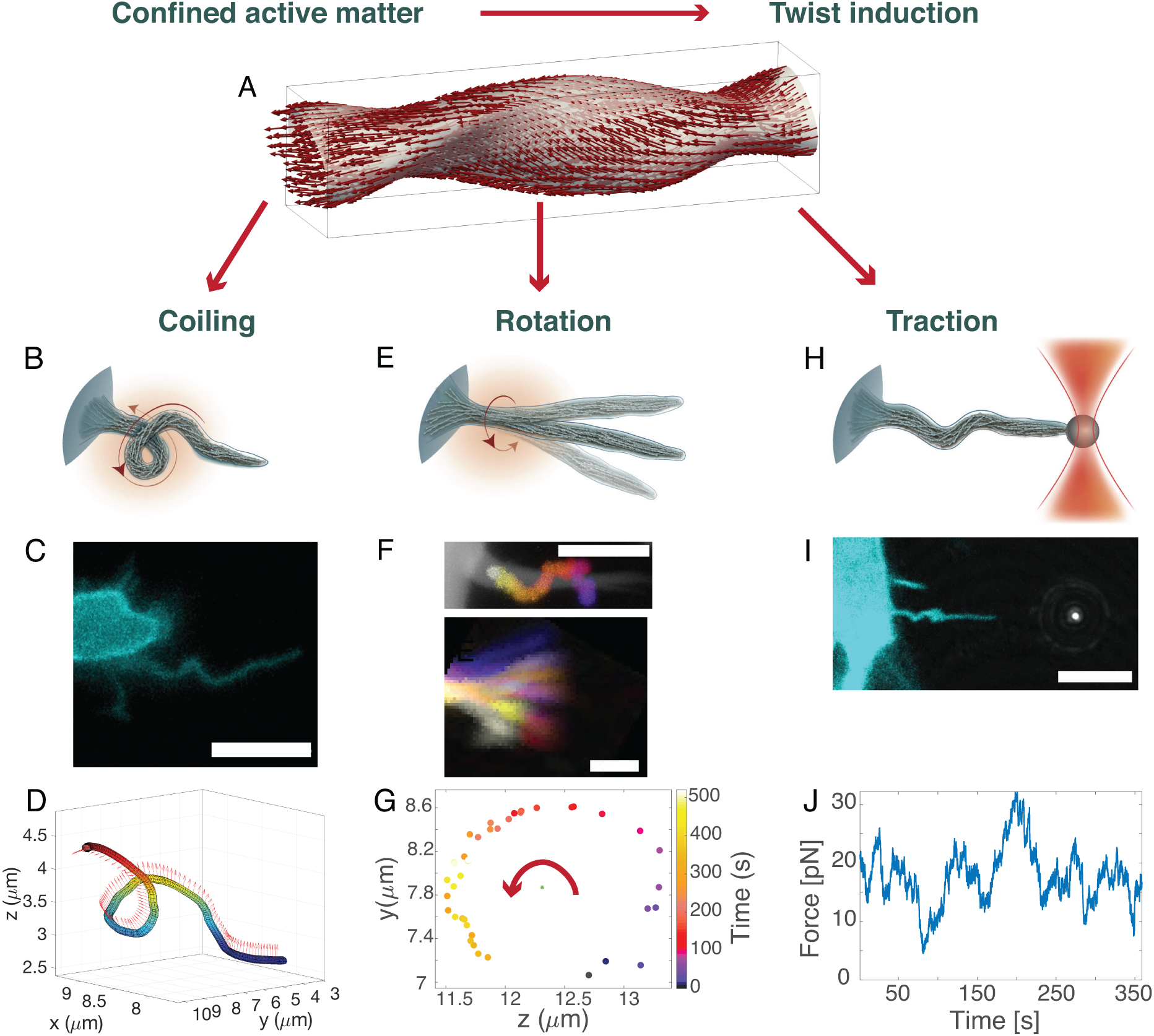
Twisting of the active filamentous actin core of filopodia is responsible for coiling, rotation, and traction generation of filopodia. (A) Filopodia twist is induced by active filaments under confinement of the filopodia membrane. (B-D) Coiling shown by a schematic depiction (B), fluorescent image, scale bar is 5 *μ*m (C), and 3D tracing of a filopodium from a MCF7 cell (cyan, EGFP Lifeact-7) (see also Figure S4B) (D). Arrows in indicate the binormals at different positions along the curve. (E) Schematic depiction of a rotating free filopodium. (F) Top: Confocal image of a filopodium extended from a MCF7 cell (white, EGFP Lifeact-7) overlayed with time color-coded images of the bead rotating around the actin shaft. Scale bar is 5 *μ*m. A vitronectin coated bead, indirectly attached to the actin fibers via transmembrane integrins, performs a spiral motion around the filopodium towards the cell body as a result of the internal twisting of actin shaft and retrograde actin flow (see also Figures 2, S2a). Bottom: Gabor-filtered *z*-projection of the movement of a filopodial tip of an EGFP Lifeact-7 labelled MCF7 cell at consecutive color-coded time points. Scale bar is 1 *μ*m. (See Figure 3). Actin shaft twisting and tip rotation occur at similar rates. (G) Tip tracking in the *yz* plane reveals counterclockwise rotation as seen from the tip towards the cell body. (H) Schematic depiction of an optically trapped filopodium performing traction due to buckling and thus apparent shortening induced by accumulation of twist. (I) Fluorescent image of an optically trapped and buckling filopodium extracted from a HEK293T cell (cyan, EGFP Lifeact-7) using a 4.95 *μ*m bead. Scale bar is 5 *μ*m. (J) Force profile from a filopodium extracted from a mESC in S/L medium.

The rotational behavior of filopodia has been largely unexplored in literature. However, the periodic sweeping motion of macrophage filopodia was found to be 1.2 rad/s (0.191 Hz) as measured in two dimensions. ^22^ Growth cone filopodia were also tracked in two dimensions orthogonal to the axis of the filopodium and the frequency was found to be ∼ 0.016 Hz.^6^ In Ref.^17^ the actin structure of filopodia was imaged and small buckles were observed to rotate around the actin shaft, thus strongly indicating that actin has the ability to spin within the filopodium.

We explicitly performed 3D tracking of the whole filopodium and its tip, see Figure 7B-D, and made complimentary measurements of the spinning of the actin shaft within the filopodium, see Figure 7E-F. The results showed that the actin shaft has the ability to spin with a similar frequency as the circular movement of the whole filopodium. These results strongly suggest that the spinning of the actin together with filopodia bending and growth, is responsible for the 3D motion and buckling of filopodia.

When a filopodium is free, this gives rise to the tip rotation with a frequency of (0.0084± 0.0084) Hz. In the presence of friction arising from being embedded in a 3D matrix we observe that rotation slows down and filopodia rotate with a frequency of (0.0024 ± 0.0011) Hz. The external resistance to rotation experienced by filopodia leads to accumulation of twist and hence to helical buckling of the filopodium (Figure 7B-D). This mechanism can be explained by twisting a rubber cable with one hand while holding the other end tight. The cable will accumulate twist, start to buckle and coil into an helical shape as shown in supplementary Figure S8. During buckling and coiling such a structure will shorten and therefore generate a pulling force.

When actin is not specifically linked to an external structure which can prevent spinning of the actin, buckling can still occur. In the presence of friction between the membrane and the actin, for example from the transmembrane proteins such as integrins and peripherally binding proteins like I-BARs,^53^ torsional twist can accumulate in the actin shaft and cause buckling.^17,54^ Buckles form when twist is released and converted into bending energy within the actin shaft. This leads to an apparent shortening of the filopodium as observed when holding the filopodium by an optical tweezer^17^ and thus contributes to a traction force in addition to the force arising from the retrograde flow of actin^18^ (Figure 7H-J).

We found the filopodia to predominantly rotate in a counterclockwise orientation, as seen from tip the towards the cell body. Of all resolvable rotations we found 46 % of the filopodia to rotate in a counterclockwise orientation while 7 % rotated in a clockwise orientation as seen from the tip towards the cell. The orientation of 47 % of the filopodia rotations could not clearly be resolved due to convoluted lateral movements and rotations. Interestingly, a similar sweeping motion was also measured for the movement of growth cone filopodia in ref.^6^ which could indicate that filopodia are generically proned to exhibit counterclockwise rotation when observed from the perspective of the tip. However, to properly resolve the orientation of the spinning actin shaft it is necessary to perform experiments as in Figure 2 where the actual spinning of the actin shaft can be isolated from the lateral movement of the filopodia.

Twist in actin bundles emerges naturally when active filaments are confined in channels with dimensions similar to a filopodia tube as shown by active matter simulations, see Figure 7A. The induced chirality is random, but can be biased by active torque dipoles in the system. Such dipoles indeed exist in natural filopodia in the form of myosins exhibiting chiral motion along actin bundles as demonstrated for the filopodia associated motors myosin V^24^ and myosin X.^51^ Both of these motors walk towards the tip in a filopodium and spiral around the actin in a clockwise orientation as seen from the tip thus inducing counterclockwise chirality as seen from the perspective of the tip. Our experimental data indeed show a preference for counterclockwise orientation of the twist, but the presence of both twist orientations supports the theoretical predictions of a hydrodynamic instability being the origin of the twist.

To exclude possible artifacts arising from the transiently expressed fluorescent actin we also imaged cells which had the actin labelled with a membrane permeable probe which binds to F-actin in living cells (SiR actin). These cells showed similar filopodial activity as we observed in cells transfected with EGFP Lifeact-7, see SI movies S3b,c.

The observation of similar filopodial dynamics in early stem cells confirms the generic nature of filopodial dynamics. The state of stem cells used here resemble the inner cell mass (ICM) cells, but their state can be controlled by different culture media. Cells cultured in serum-free medium supplemented with leukemia inhibitor factor (LIF) and small molecule inhibitors GSK3 and MEK (2i medium), genetically represent cells from the ICM of mouse blastocyst corresponding to 3.5 days post fertilization and are highly pluripotent.^55,56^ Cells grown in Serum-LIF (S/L) medium are in a primed state towards epiblast.^56^ The colonies of cells grown in these two media exhibit remarkably different morphologies; cells cultured in S/L media are more spread out (Figure S5A (D)) and form a monolayer of cells while 2i cultured cells grow together into an embryonic body (Figure S5A (C)). The general role of filopodia in early development is less clear. Their presence has been detected in both the process of embryonic compaction, where they were shown to sustain tension, which was concluded form images, but the force was not quantified.^9,57^ Our quantitative data show that naive pluripotent embryonic stem cells are able to deliver a traction force on the order of 10 pN for each filopodium. However, whether the spinning motion of actin does occur during compaction, where the filopodia are penetrating other cells, remains an open question. Filopodia in mesenchymal stem cells have been shown to function as rails facilitating transport of morphogens between cells.^2^ Such tunnelling tethers or cytonemes have been observed in many cell types and we have also detected rotation of such structures in Hepa 1-6 mouse hepatocytes (SI movies 3b,c), however, the functional role of the rotation remains elusive in these structures.

## Conclusion

Filopodia are extremely versatile surface structures with ability to explore the 3D extracellular space by simultaneous growth, bending, and rotary motion. We find that the actin shaft inside filopodia generally spins and is responsible for the circular motion of filopodia which allows them to explore their 3D extracellular environment. Using a hydrodynamic model of actin filaments we show that the spinning can originate from symmetry breaking and subsequent twist generation by active filaments under a confinement with similar dimensions as the filopodia tube. Signatures of twist are observed experimentally as helical buckles and coils in a variety of cell lines and can give rise to shortening of the actin shaft and therefore generation of a traction force. Additionally, the functional implication of twist generation in filopodia for cellular processes could be to allow the cell to explore their 3D extracellular environment, to navigate through dense networks of the ECM and to assist in chemical sensing.

## Supporting information

collagen I bending arr.avi} Bending filopodia from the KPC cell (cyan, EGFP Lifeact-7) from Figure 1C,D and Figure S4a in 4 mg/ml collagen I (gray, re

Filopodia on a mESC (cyan, EGFP Lifeact-7) grown in 2i medium.

Bending filopodia from KPC cell (cyan, EGFP Lifeact-7) in 4 mg/ml collagen I (gray, reflection) over a time span of 399s. Red arrows highlight bending

## Acknowledgements

PMB and NL acknowledge support from the Danish Council for Independent Research, Natural Sciences (DFF-4181-00196) and LBO acknowledges support from the Danish National Research Foundation (DNRF116). A. D. acknowledges support from the Novo Nordisk Foundation (grant No. NNF18SA0035142), Villum Fonden (Grant no. 29476), Danish Council for Independent Research, Natural Sciences (DFF-117155-1001), and funding from the European Union’s Horizon 2020 research and innovation program under the MarieSklodowskaCurie grant agreement No. 847523 (INTERACTIONS). J.T.E. is funded by a Hallas Møller Stipend from the Novo Nordisk Foundation. We thank Joshua Brickmann for providing us with embryonic stem cells, Anne Benedicte Mengel Pers for providing hepatocytes and Agnieszka Kawska for help with schematic illustrations.

## 1 Materials and Methods

### 1.1 Cell culture

All cells were grown in vented T25 flasks (BD Falcon) in a sterile environment at 37°C in a humidified 5 % CO_2_ incubator. The cells were passaged at around 80 % confluence.

#### HEK293, HEK293T, and MCF7

Cells were grown in DMEM growth medium ([+] 4.5 g/L D-Glucose, [+] L-Glutamine, [+] 110 mg/L Sodium Pyruvate, Gibco, supplemented with 10 % FBS, 1 % PenStrep) and passaged by washing with 2 ml DPBS (1X, [-] CaCl_2_, [-] MgCl_2_, Gibco or PBS pH 7.4, Gibco), detached using 1 ml TrypLE Express ([-] Phenol red, Gibco) and resuspended in growth medium.

#### MCF7-p95ErbB2

MCF7-p95ErbB2 cells described previously^1^ were grown in DMEM growth medium ([+] 4.5 g/L D-Glucose, [+] L-Glutamine, [+] 110 mg/L Sodium Pyruvate, Gibco, supplemented with 10 % FBS, 1 % PenStrep, further supplemented with 200 *μ*g/ml G418, 1 *μ*g/ml Puromycin, 0.5 *μ*g/ml tetracycline). To induce p95ErbB2 expression, the cells were trypsinized and washed 4-5 times in 20 ml of PBS (+CaCl_2_) to remove all tetracycline and then plated in a new tissue culture flask. When cells were induced, they were keep in DMEM growth medium ([+] 4.5 g/L D-Glucose, [+] L-Glutamine, [+] 110 mg/L Sodium Pyruvate, Gibco, supplemented with 10 % FBS, 1 % PenStrep). The “spiderlike” phenotype of the MCF7-p95ErbB2 cells appears after 3-5 days.

Uninduced MCF7-p95ErbB2 cells, meaning cells grown in growth medium containing tetracyline, will express some ErbB2 as the promotor is leaky, but will resemble more normal MCF7 cells in their activity levels.

#### KPC, KP^fl^C

Cells were cultured in DMEM containing GlutaMAX, 10 % FBS, and 1 % PenStrep.

#### Stem Cell Lines

Mouse embryonic stem cell lines (mESCs) HV.5.1, HV.10.6, and EO.1 were obtained from the Brickman Group (DanStem, University of Copenhagen, Denmark). HV.5.1 cells are equipped with a Venus reporter which represents the expression state of Hex, line. HV.10.6 expresses an mCherry labeled variant of this reporter and a fluorescently tagged H2B protein, which is one of the main histone proteins. EO.1 cells express an EGFP E-cadherin fusion protein and an mCherry labeled reporter for the pluripotency factor Oct4. HV.5.1 had been passaged 23 times prior to our experiments, HV.10.6 26 times and EO.1 12 times, respectively. The cultures were sustained for no longer than 25 subsequent passages.

Stem cells were either cultured in Serum/LIF (S/L) or 2i culture medium. *Serum/LIF (S/L) medium*. The addition of the cytokine Leukemia Inhibitory Factor (LIF) maintains a pluripotent state of the stem cells.^2–4^ Cells were cultured in Glasgow modified Eagle’s medium (GMEM, Sigma Aldrich, Germany) supplied with 5.5 ml non-essential amino acids (Gibco Science, Paisley, UK), 5.5 ml glutamine, 5.5 ml sodium pyruvate (Gibco Science, Paisley, UK), 560 *μ*l 0.1 mM 2-merceaptoethanol (Sigma-Aldrich, St. Louis, MO, USA), 50 ml fetal bovine serum (FBS) (obtained from DanStem, University of Copenhagen, Denmark), 560 *μ*l 1000 U/ml LIF (obtained from DanStem, Copenhagen University, Denmark) per 500 ml of medium. The entire pile of FBS was bought from the manufacturer by DanStem to ensure consistency in the culture medium. For S/L culture, the medium was aspirated, and rinsed once with DPBS and detached with 0.025 % trypsin in DPBS, incubated for 4 min at 37° C and suspended in 5 ml of S/L medium. The cells were then centrifuged for 3 min at 1890 rpm and re-suspended in 5 ml of S/L.

*2i medium*. In 2i medium, cells remain in a more primitive state. 2i composition: 1:1 mixture of DMEM (Gibco Science, Paisly, UK) and F12 medium, modified N2 medium (25 *μ*g/ml insulin, 100 *μ*g/ml apo-transferrin, 6 ng/ml progesterone, 16 *μ*g/ml putrescine, 30 nM sodium selenite, obtained from DanStem, University of Copenhagen, Demark), 50 *μ*g/ml bovine serum albumin fraction V, combined with a 1:1 mixture of Neurobasal medium supplemented with B27 (Gibco, Paisley, UK). The culture medium contained two cytokines, a GSK3 inhibitor with a final concentration of 1 *μ*M and ERK inhibitor with a final concentration of 3 *μ*M. The cells were cultured in 50 ml Falcon flasks (Falcon Science, Paisley, UK) coated with EmbryoMax ultrapure water with 0.1 % gelatin (Merck Millipore, Billerica, MA, USA) for 10 minutes. At a confluence of approximately 90 % the cells were harvested. Cells detach from the culture flask with time. To retain a large number of cells, the culture medium was collected in a 15 ml Falcon tube and centrifuged for 3 minutes at a speed of 1890 rpm prior to trypsinization. The supernatant was aspirated and the pellet was re-suspended in 1 mL of 0.025 % trypsin by pipetting. The suspension was added to the flask in which the cells had been cultured to detach the clusters of cells that had remained bound to the substrate and incubated at 37°C for 4 min. Cells were transferred to a 15 ml Falcon tube and 5 ml of N2B27 was added to reduce the usage of the cytokines and LIF. The suspension was centrifuged with the same settings and re-suspended in 5 ml of N2B27 medium, by pipetting carefully up and down. Finally, a volume of the cell suspension was added to a new culture flask together with 5 ml of fresh 2i medium.

### 1.2 F-actin labelling

#### Transient transfection of HEK293, HEK293T and MCF7 cells

The day prior to transfection, the cells were seeded on a 24 well cell culture plate (Orange Scientific). On day of transfection, the medium in each well was removed and exchanged with 500 *μ*l fresh growth medium per well. Then 1 *μ*l of EGFP Lifeact-7 plasmid (mEGFP-Lifeact-7 was a gift from Michael Davidson; Addgene plasmid # 54610), 100 *μ*l Opti-MEM ([+] HEPES, [+] 2.4 g/L Sodium Bicarbonate, [+] L-Glutamine, Gibco), and 1 *μ*l Lipofectamine LTX Reagent (Invitrogen) were carefully mixed in an Eppendorf tube and incubated at RT for at least 30 min. This mix was sufficient for a single well. Usually, the cells in at least 3 wells were transfected at a time. The next day, the cells were detached from the wells as described above, the contents of the 3 wells were mixed together. 100 *μ*l of those cells and ca 1 ml of growth medium was added to a glass bottom dish (35 mm, No. 1.5 coverslip, 20 mm glass diameter, uncoated, MatTek) and left to adhere and express for another day in the incubator. The samples for experiments were then used 1 or 2 days after transfection.

#### Transient transfection of KPC and KP^fl^C

KPC and KP^fl^C cells were transfected with mEmerald LifeAct-7 (a kind gift from the Ivaska lab, Turku Centre for Biotechnology) using Lipofectamine 2000 as described above. After an incubation period of 12 h they were embedded in collagen I matrices.

#### Transient transfection of mESCs

Cells were transfected with EGFP LifeAct-7 or RFP LifeAct-7 using Lipofectamine P3000 in 24 well plates. The day before transfection, the cells were plated at high density (70 − 90% confluency). On the day of transfection, a transfection mixture containing Opti-MEM, plasmid, Lipofectamine P3000, P3000 reagent, and medium was mixed and incubated for 12 minutes at room temperature. The medium was aspirated from the cells and the mix was subsequently added and was equally distributed over the well. Thereafter the well was filled with fresh medium. Although the manufacturer recommends to use Opti-MEM medium for better transfection efficiency, the cells were cultured in S/L or 2i medium, respectively, to maintain an optimal pluri- or multipotent state. The cells were used for experiments between 24 and 48 hours after transfection.

#### SiR actin labeling

Hepa 1-6 cells were incubated in SiR actin (SiR actin kit, Spirochrome, 1 : 2000) and Verapamil (1 : 1000) for 5 h.

### 1.3 Collagen I gels

Collagen mixtures of 4 mg/ml were prepared by mixing high concentration acid-extracted and cross-linked rat tail collagen I, sterile phosphate-buffered saline (PBS), and 5X collagen buffer containing 0.1 M HEPES, 2 % NaHCO_3_, and *α*-MEM. Cells were then suspended in the collagen mixtures. After polymerization, the gels were washed once and incubated with normal culture medium for 24 h.

### 1.4 Vitronectin coated beads

We prepared two stock solutions: ‘tracer beads’ with *d* = 0.989 *μ*m streptavidin coated Flash Red fluorescent beads (Bangs Laboratories) and ‘tether beads’ with 4.95 *μ*m streptavidin coated polymer particles (Bangs Laboratories) coated with biotin labeled human multimeric vitronectin (VN) (Innovative Research) in PBS (pH 7.4, Gibco) which were stored at 4° C. For coating, 5 *μ*l of beads were washed twice in 1 ml of PBS. After the second washing step, the beads were re-suspended in 50 *μ*l of PBS containing 2 *μ*l of VN, enough to exceed the maximal binding affinity of the streptavidin coated beads. The beads were incubated for 20 min at a shaking platform (Eppendorf MixMate, Hamburg, Germany) with a speed of 450 rpm. To remove unbound VN, the beads were centrifuged at 12000 rpm for 5 min and re-suspended in 1 ml of PBS.

### 1.5 Imaging sample preparation

On day of experiments, 400 *μ*l of ‘imaging DMEM’ ([+] 4.5 g/L D-Glucose, [+] L-Glutamine, [+] 25 mM HEPES, [-] Sodium Pyruvate, Gibco) was added to an Eppendorf tube together with 3 *μ*l and 7 *μ*l of 1 *μ*m and 4.95 *μ*m VN coated beads, respectively. The medium in a MatTek dish with transfected cells (see section ‘Transient transfection’) was carefully removed and replaced with the described bead mix in imaging DMEM.

## 2 Experimental setup and procedures

### 2.1 Confocal microscopes with integrated optical trap

Experiments were conducted on two different setups. One setup consists of an inverted Leica TCS SP5 II confocal microscope with a 63X water objective (1.2 NA, Leica) with an integrated optical trap based on a 1064 nm laser (Nd:YVO4, 5 W Spectra Physics BL106C, *λ* = 1064 nm, TEM∞) and a photodiode detection system. The laser beam was tightly focused by the water-immersion objective and the trapping laser light was collected by a condenser (Leica, P1 1.40 oil S1) located in the back-focal plane and focused onto a quadrant photodiode (S5981; Hamamatsu). A three-dimensional LabVIEW controlled piezoelectric stage (Nano-Drive, MCL) allowed positioning of the sample relative to the laser focus with nanometer precision. For some experiments, the sample stage was heated to 39° C using a stage heater which, due to thermal losses, results in a sample temperature of about 37° C inside the sample. Data were acquired by an acquisition card (NI PCI-6040E) at a sampling frequency of 1 kHz (for force measurements) and 22 kHz (for force calibration) and processed by custom-written LabVIEW programs (LabVIEW 2010; National Instruments).

The second setup, mostly used for force measurements and stem cell experiments, is described in.^5^

### 2.2 Free filopodia and filopodia in collagen rotations

Cells in MatTek dishes containing imaging medium or grown in collagen I gels were placed on the confocal microscope and filopodia movement was imaged via *z*-stacks over time. The mEmerald Lifeact-7 or EGFP Lifeact-7 expressing cells were excited at 488 nm and the collagen I gel was imaged in reflection mode using the 633 nm laser without enhanced dynamics.

### 2.3 Bead rotations on tether

For experiments where the movement of a VN coated bead along a tether was followed, a MatTek dish with transfected cells and VN coated beads was placed on the sample stage. In the following, we distinguish two types of VN coated beads: ‘tether beads’ with *d* = 4.95 *μ*m, and ‘tracer beads’, with *d* = 0.989 *μ*m, see section 1.4 for VN coating details. During tether extraction we imaged the sample in bright field mode to avoid confocal laser radiation before the actual data acquisition. A tether bead was optically trapped and brought close to a (usually isolated and spread out) cell. The bead was carefully pressed against the cell membrane to allow attachment and then slowly moved a distance of about 10 *μ*m (using the LabVIEW controlled piezo stage) such that a membrane tether was extracted between cell and bead. Once a tether was extracted, we switched to confocal mode. The EGFP Lifeact-7 expressing cells were excited at 488 nm, the flash red tracer beads at 633 nm. The tether bead was carefully pushed onto the sample glass surface such that it got stuck and the trapping laser could be turned off. A control *z*-stack was acquired (the tether beads were imaged in reflection mode using the 633 nm laser without enhanced dynamics) to ensure the tether was held and did not stick on the glass surface. Then a tracer bead was caught using the optical trap and carefully attached onto the tether close to the tether bead. Confocal *z*-stacks over time were acquired to follow the tracer bead’s rotation around the tether towards the cell body.

## 3 Data analysis

Image processing and analysis were done using ImageJ and custom written MATLAB scripts.

### 3.1 Tracking the movement of a bead attached to a filopodium

To localize the position of an attached bead on a filopodium, the *xy* position of the center of the bead and the filopodium were extracted from the *z*-projection of the image stack, see Figure S6. Afterwards, the *z*-positions in each time point were extracted from the orthogonal view of the stack, shown in Figure S6B, which corresponds to the red dashed line in Figure S6A.

### 3.2 Tracking the rotation of filopodial tips

To quantify the tip rotation of filopodia, we increased the resolution of the images using a Gabor filter.^6,7^ Then the center of a filopodium was extracted by fitting Gaussian function to the orthogonal view along the filopodium, see Figure S7.

### 3.3 Tracking helical filopodial coils

We used a 3D segmentation plugin software (Simple Neurite Tracer) in ImageJ^8^ to track filopodia after removing the background intensity in images using Gabor and Gaussian blur filters (Figure S8).

## Simulation Methods

We use the 2D equations of active nematohydrodynamics based on the theory of liquid crystals, which have proven successful in describing spatio-temporal dynamics of the cell cytoskleton, including acto-myosin complexes^9–12^ and microtuble bundles powered by kinesin motor proteins.^13–15^ The orientational order of microscopic actin filaments is represented by the nematic tensor 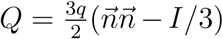, with *q* the magnitude of the orientational order, *n* the director, and *I* the identity tensor, which evolves as

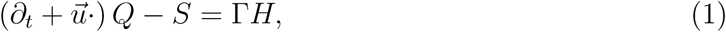

where Γ is a rotational diffusivity and the co-rotation term

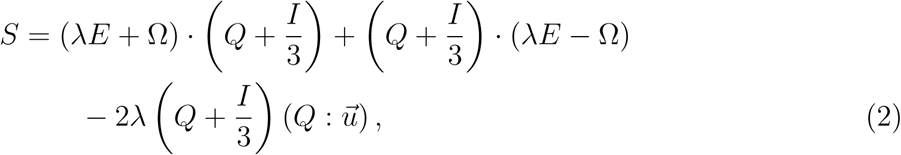

accounts for the response of the orientation field to the extensional and rotational components of the velocity gradients characterised by the strain rate 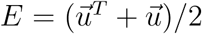 and vorticity 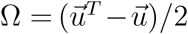 tensors. The relative strength of extensional and rotational flows is determined by the alignment parameter *λ*. Therefore, the alignment parameter accounts for the different responses of particles of different shapes to the symmetric and asymmetric parts of the velocity gradient tensor.^16^ Mapping the alignment parameter to the Leslie–Ericksen equation for liquid crystal dynamics gives 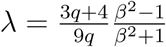,^17^ where *β* = *a/b* is the ratio of the length of the cell along its axis of symmetry, *a*, to its length perpendicular to this axis, *b*. Therefore, for prolate ellipsoids *β >* 1, while for oblate ellipsoids *β <* 1 and for spherical particles *β* = 1, which correspond to *λ >* 0, *λ <* 0, and *λ* = 0, respectively. Actin filaments are rod-shaped elongated particles and therefore are characterized by *λ >* 0. The relaxation of the orientational order is determined by the molecular field,

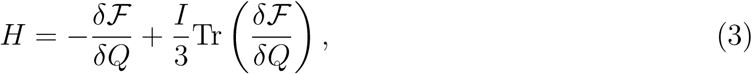

where ℱ = ℱ_*b*_ + ℱ_*el*_ denotes the free energy. We use the Landau-de Gennes bulk free energy for ℱ_*b*_,^16^

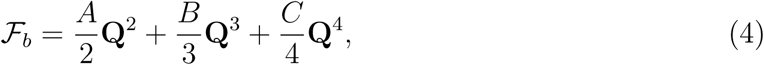

and 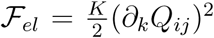 describes the cost of spatial inhomogeneities in the order parameter, assuming a single elastic constant *K*. There are in general three elastic constants, corresponding to bend *K*_*b*_, splay *K*_*s*_, and twist *K*_*t*_ in 3D systems. However, setting different values of *K*_*b*_, *K*_*s*_, and *K*_*t*_ does not change the mechanism of the hydrodynamic instability.

The velocity field is evolved according to the incompressible Navier–Stokes equation:

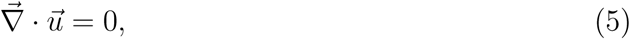

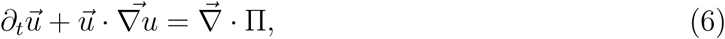

which reduces to the force balance equation 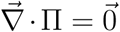 in the low Reynolds number limit relevant to actin filaments mechanics, with a stress tensor **Π** that must account for contributions from the viscous stress Π^visc^ = 2*ηE* and the elastic stress

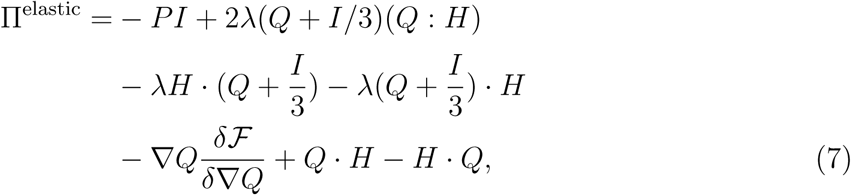

which includes the pressure *P*.^18^ The active contribution to the stress is accounted for by Π^act^ = −*ζQ*,^19^ such that any gradient in **Q** generates a flow field, with strength determined by the activity coefficient, *ζ*. This active stress term accounts for the local stresses generated by active processes in the cells, including actomyosin polymerization and contractility.^9,10^

The equations of active nematohydrodynamic are solved using a hybrid lattice Boltzmann and finite difference method^20^ that does not include thermal fluctuations. The momentum equation is solved using the lattice Boltzmann method to resolve the hydrodynamics, and the method of lines is implemented to determine the order paramter in Eq. 1. A finite-difference approach is used for spatial discretisation of Eq. 1 and the temporal evolution is obtained through an Euler integration scheme.

A three dimensional rectangular domain with a square cross-section is used for simulations. The length (*L*) of the channel was fixed as 128 while the height (*h*) was varied between 4 to 32. In our study, simulations are initialized with a stagnant fluid and randomly oriented director field. Zero velocity and no flux of **Q** at the walls are used as the boundary conditions, therefore no anchoring boundary condition is imposed for the orientational order parameter on the channel walls.

The parameters used in the simulations are *A* = 1.0, Γ = 0.34, *η* = 2*/*3, and *λ* = 0.7, in lattice units. As usual in lattice Boltzmann schemes, discrete space and time steps are chosen as unity and all quantities can be converted to physical units in a material dependent manner.^20–22^ In order to map the simulation parameters to physical units we consider the typical length of the filopodia as the characteristic length *L*_0_ ∼ 10 *μ*m, and the experimentally measured viscosity and elasticity of the actin/myosin mixtures^11,12^ as the characteristic viscosity *η*_0_ = 0.1Pa.*s* and force units *F*_0_ = 10 pN, respectively. This maps the confinement sizes studies here to the range ∼ 0.2 − 2 *μ*m, elasticities to the range ∼ 0.1 − 1.0 pN, and the activities to ∼ 0.01 − 0.1Pa, consistent with the values reported for *in vitro* two-dimensional extract of actin/myosin mixtures.^12^

Furthermore, the magnitude of the torque dipole can be estimated based on analytical derivation of,^23^ which suggests:

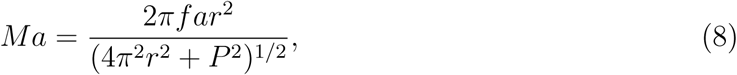

where *M* is the torque monopole, *a* ∼ 100 nm distance between filaments, *f* ∼ 10 pN is the force generated by myosin motors, *r* ∼ 5 nm the radius of actin filament, and *P* ∼ 72 nm is the helical pitch of the actin filaments. Using these values the ratio of the active stress associated with the torque dipole *ζ*′ to the active stress due to force dipole *ζ* can be estimated as *ζ*′*/ζ* ∼ *M/fa* ∼ 0.1, which is the value mentioned in the main text. Therefore, the mechanism of symmetry-breaking and twist generation is governed by hydrodynamic instabilities that are induced by the active stress associated with force dipoles. The role of torque dipole is to set the direction of the twist: in the absence of a torque dipole *ζ*′ = 0 (results in the main text) clockwise and counter-clockwise rotations are equally likely. Including a torque dipole is expected to bias this twist towards clockwise and counter-clockwise rotation depending on the sign of *ζ*′. To test this, we performed simulations with *ζ*′ = (0, 0.1 × *ζ, ζ*) and compared the distribution of clockwise and counter-clockwise rotations for 20 simulations - for each case - with random initial conditions. As evident from Figure S9 increasing the value of *ζ*′ clearly increases the biased rotation and when it becomes comparable to the value of active stress associated with force dipoles it can deterministically set the direction of rotation.

## 4 Additional Figures

**Fig. S1.**
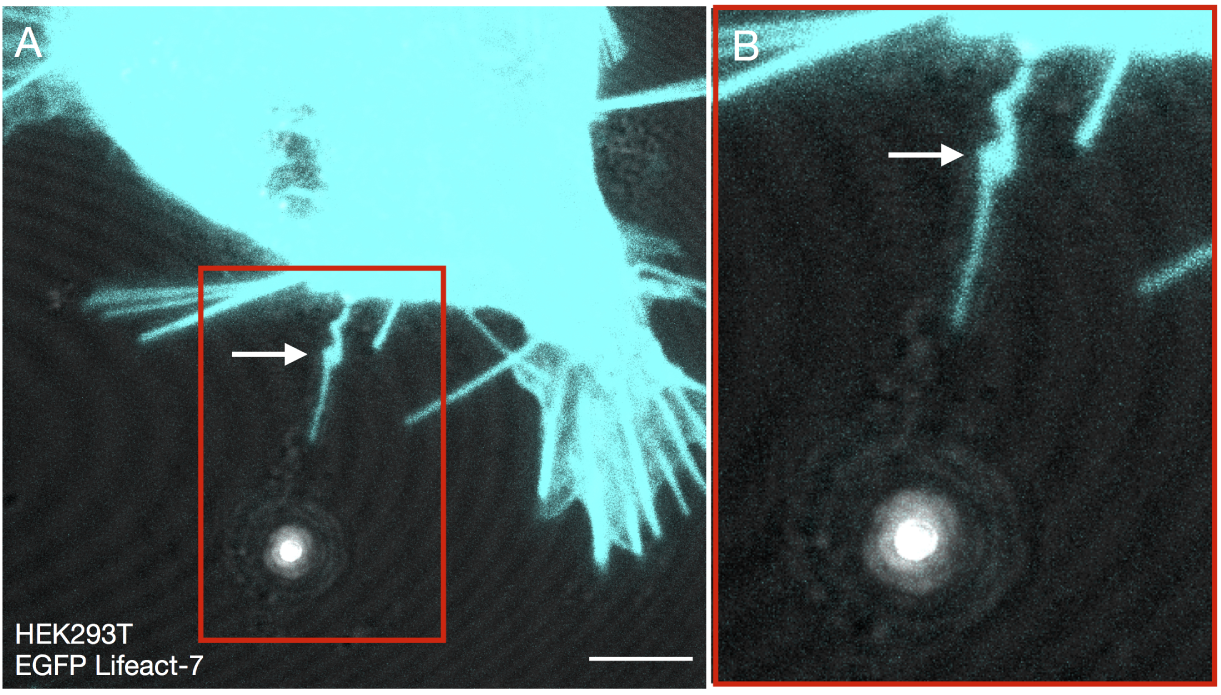
Filopodia rotation is not an artifact of confocal laser illumination. (A) Confocal *z*-projection of tether extracted from a HEK293T cell (cyan, EGFP Lifeact-7) using a 4.96 *μ*m bead shows a buckle (white arrow) stemming from rotation of the filopodium. The tether was extracted in bright field mode without exposing the cell to laser illumination prior to confocal imaging. Scale bar is 5 *μ*m. (B) Zoom in to the red marked region in (A).

**Fig. S2A.**
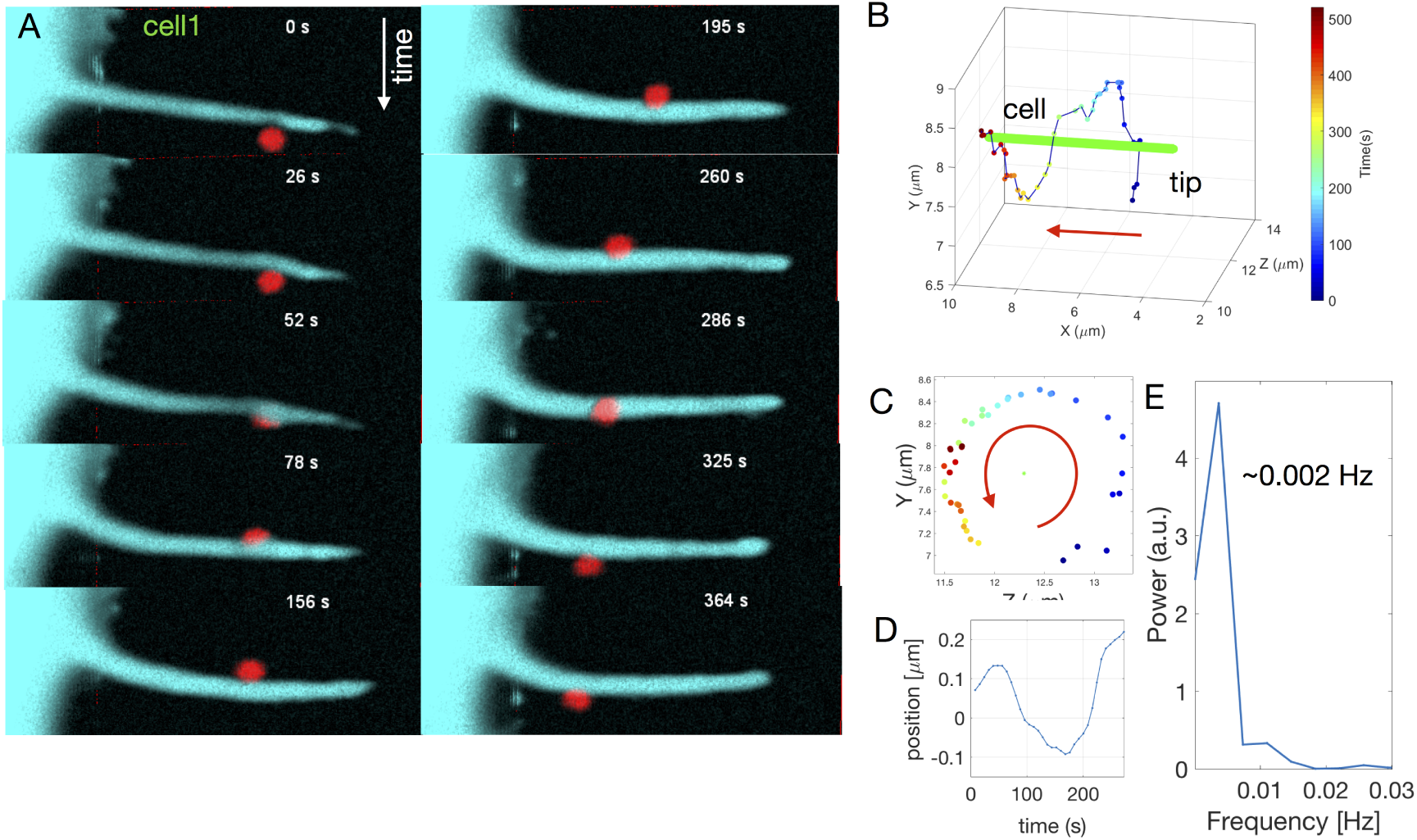
Bead rotation assay reveals that the F-actin shaft inside filopodia rotates around its own axis (additional data for cell 1 from Fig. 2B). (A) 3D reconstructed images of the rotation of a VN coated tracer bead (*d* = 0.989 *μ*m, flash red) around an extended filopodium of a HEK293T cell (cyan, EGFP Lifeact-7, cell 1 from Fig. 2B) at consecutive time points. (B) 3D trajectory of the bead in (A) (from blue to red) rotating in an counterclockwise orientation around the filopodium (green) from tip towards the cell body. (C) *xz* view of the bead movement from (A) over time, time bar is the same as in (B) shows the counterclockwise rotation (seen from tip towards the cell body). (D,E) The bead position (data from A) as a function of time (D) can be Fourier transformed to give the rotation frequency of the tracer bead 0.002 Hz.

**Fig. S2B.**
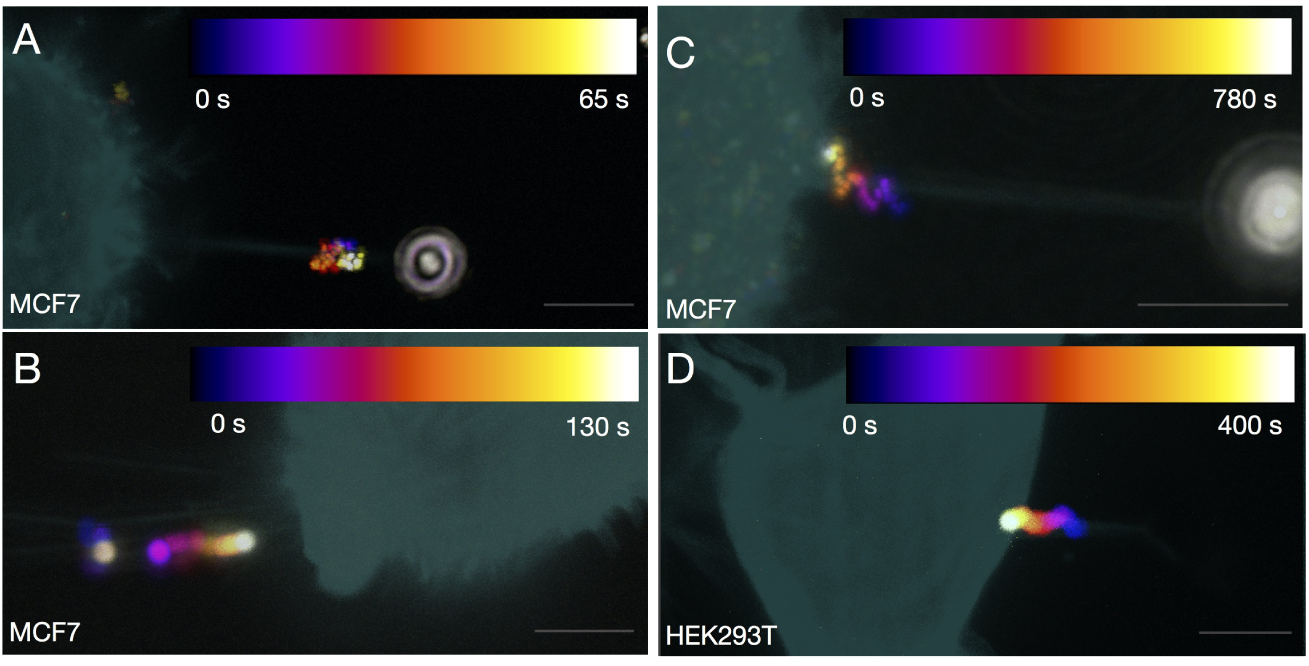
Further examples of a bead rotating around or moving along a filopodium. (A-D) Color coded *z*-projection over time of single beads rotating around (A-C) MCF7 cells (cyan, EGFP Lifeact-7) and (D) a HEK293T cell (cyan, EGFP Lifeact-7). Scale bars are 5 *μ*m.

**Table S2C.** Rotation frequencies and velocities for single tracer beads on filopodia of different cell types.

| cell type | bead rotation frequency [Hz] | bead velocity [$\mu\text{m/s}$] |
| --- | --- | --- |
| **HEK293T** | 0.002 | 0.0184 |
| **HEK293T** | 0.005 | 0.1222 |
| **MCF7** | 0.08 | 0.3676 |
| **MCF7** | 0.05 | 0.2309 |
| **MCF7** | 0.005 | 0.0312 |
| **uninduced MCF7-p95ErbB2** | 0.02 | 0.1297 |

**Fig. S3A.**
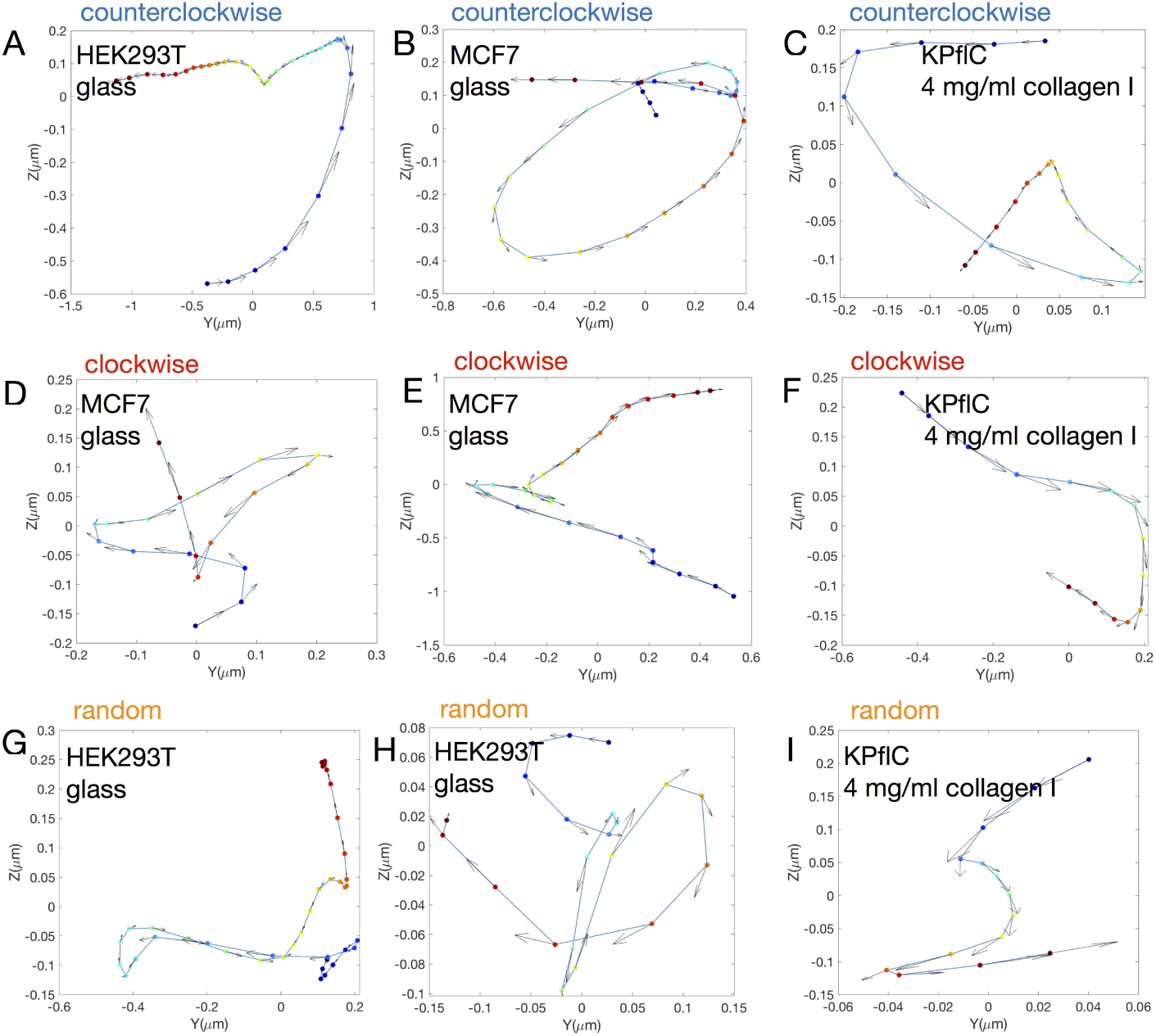
Further examples of filopodial tip rotation tracks. 46 % of tracked rotations were counterclockwise, 7 % clockwise, and 47 % random (*N* = 74). (A-C) Traces of the filopodial tip movement over time showing a counterclockwise rotation in *yz* view for the tip of a HEK293T cell on glass (A), a MCF7 cell on glass (B), and a KP^fl^C cell in 4 mg/ml collagen I (C). (D-F) Traces of the filopodial tip over time showing a clockwise rotation in *yz* view for the tip of MCF7 cells on glass (D,E), and a KP^fl^C cell in 4 mg/ml collagen I (F). (G,H) Traces of the filopodial tip over time showing random rotations in *yz* view for the tip of HEK293T cells on glass (G,H), and a KP^fl^C cell in 4 mg/ml collagen I (I).

**Table S3B.**
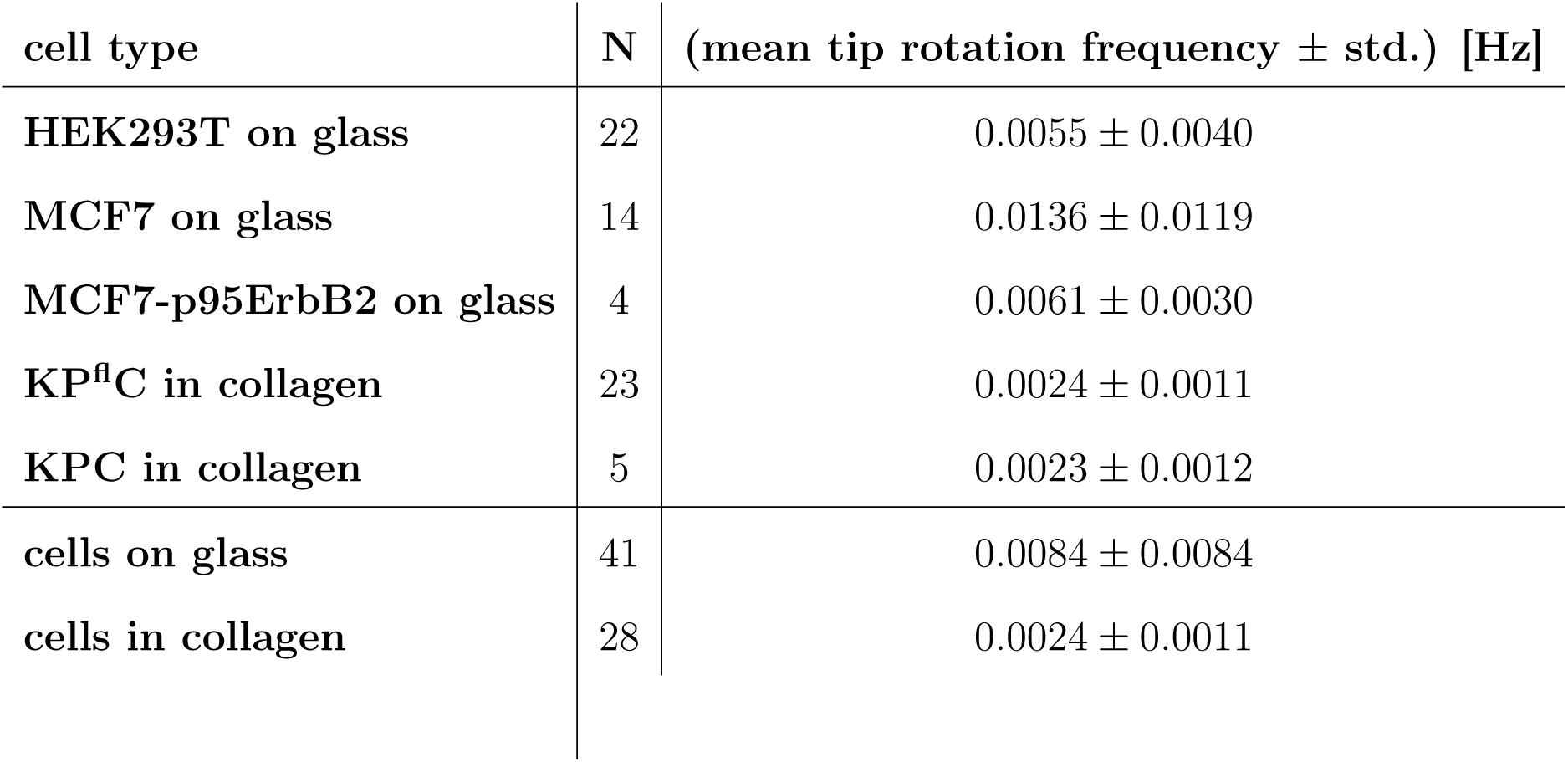
Mean filopodial tip rotation frequencies for cells on glass and embedded in collagen I gels for the data in Figure 3J.

**Table S3C.** *p*-values for the tip rotation frequency data from Figure 3J. * *p* ≤ 0.05, ** *p* ≤ 0.01, *** *p* ≤ 0.001. The *p*-values for the not mentioned couples show no significant difference between the compared populations.

| cell types | *p*-value |
| --- | --- |
| HEK293T (glass), MCF7 (glass) | 0.0058 * |
| HEK293T (glass), KP<sup>fl</sup>C (collagen) | $8.2969 \cdot 10^{-4}$ *** |
| MCF7 (glass), KP<sup>fl</sup>C (collagen) | $6.701 \cdot 10^{-5}$ *** |
| MCF7-p95ErbB2 (glass), KP<sup>fl</sup>C (collagen) | $1.0608 \cdot 10^{-4}$ *** |
| MCF7-p95ErbB2 (glass), KPC (collagen) | 0.0357 * |
| cells on glass, cells in collagen | $3.7910 \cdot 10^{-4}$ *** |

**Fig. S4A.**
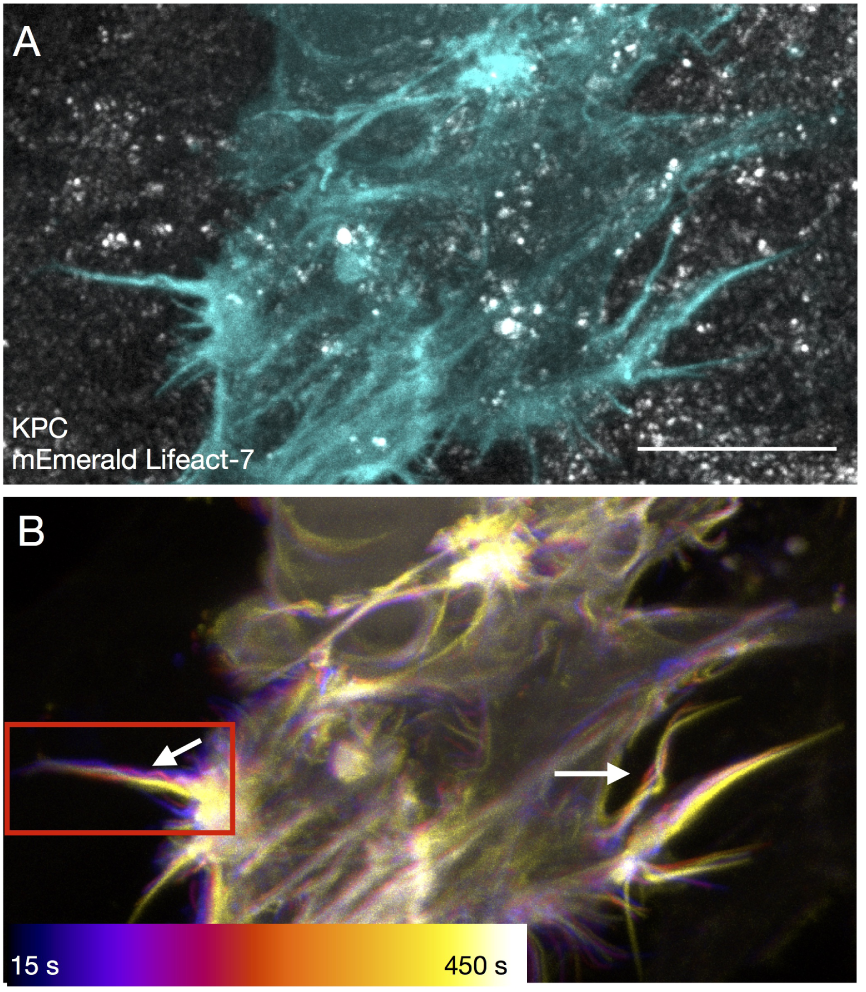
Further example of buckling filopodia from the KPC cell embedded in a 4 mg/ml collagen I gel from Fig. 1C,D. (A) Confocal *z*-projection of a KPC cell (cyan, mEmerald Lifeact-7) in 4 mg/ml collagen I gel (gray, reflection). Scale bar is 10 *μ*m. (B) Color coded temporal development of the cell from (A and Fig. 1C,D). Overlay of 6 *z*-projections acquired at different time points over an interval of 450 s. Time resolution of each *z*-stack is 15 s. The arrows highlight buckles. (B) Zoom in to the red region of (A). The right arrow marks the region shown in Fig. 1C,D.

**Fig. S4B.**
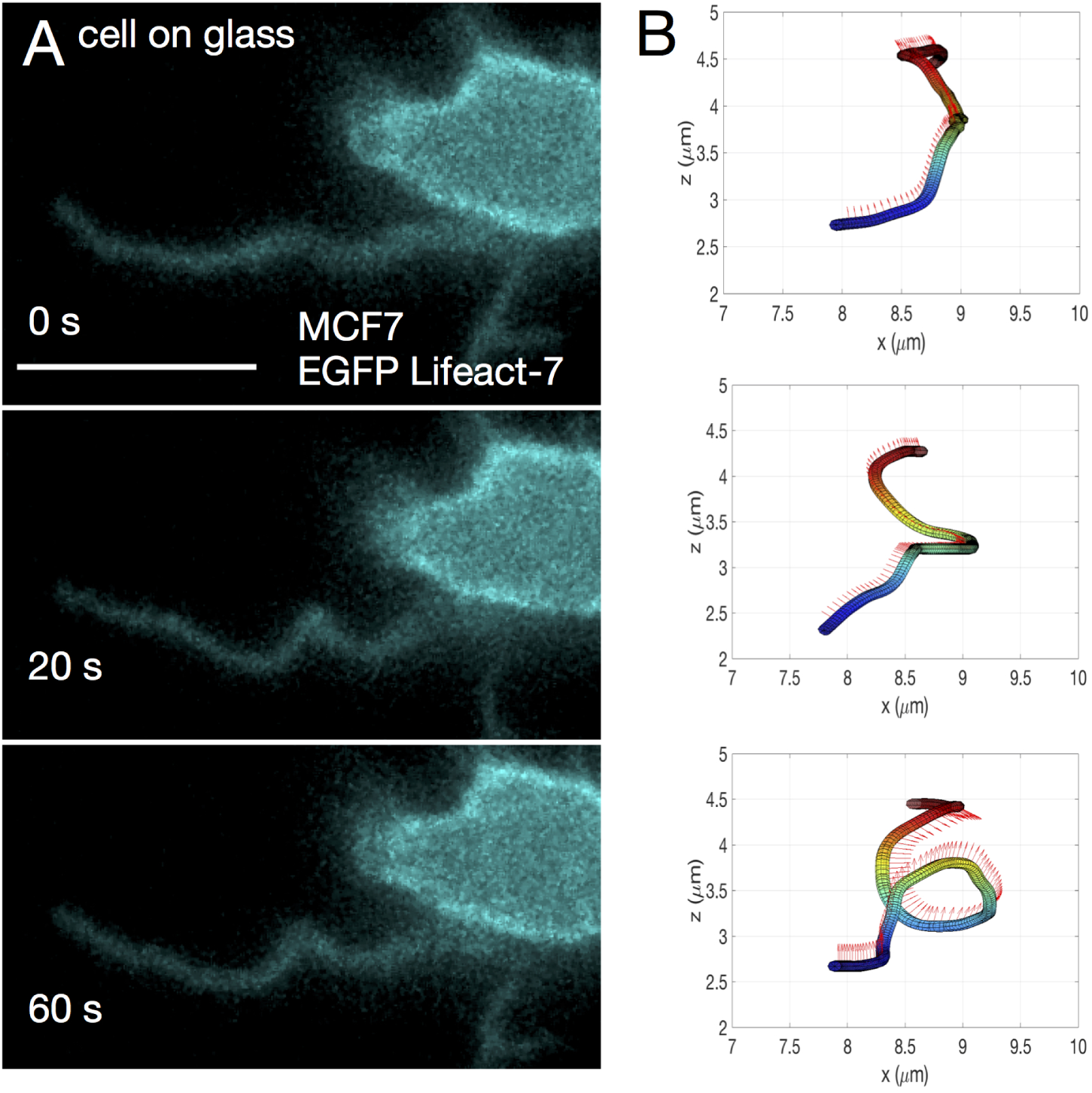
Further example of a buckling filopodium of a cell grown on glass. (A) Filopodium from a MCF7 cell (cyan, EGFP Lifeact-7) grown on glass at 3 different time points (at 0 s, 20 s and 60 s). Scale bar is 5 *μ*m. (B) 3D tracks of the filopodium corresponding to the time points from (A) show how it bends and helically coils.

**Fig. S5A.**
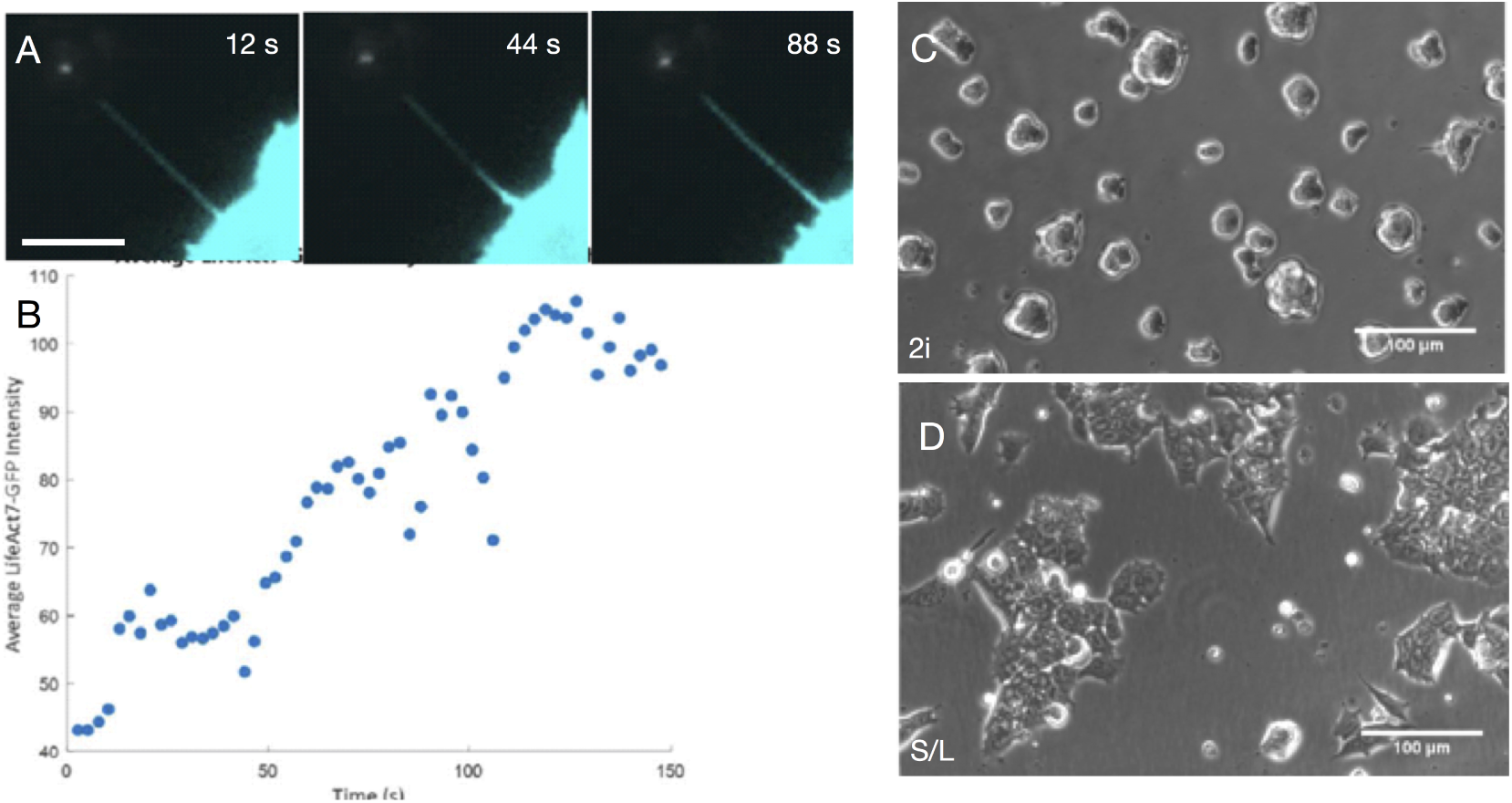
Tethers from stem cells show rich actin dynamics. (A) Tether extracted from a mES cell (cyan, EGFP Lifeact-7) grown in S/L medium at 3 different time points (12 s, 44 s, 88 s) after tether extraction. Scale bar is 5 *μ*m. (B) Quantification of the F-actin content of the tether from (A) (via the average intensity of the EGFP Lifeact-7 signal) over a time of 180 s shows how F-actin accumulates inside the tether. (C and D) mES cells cultured in 2i medium (C) are less spread out than cells grown in S/L medium (D). Scale bars are 100 *μ*m.

**Fig. S5B.**
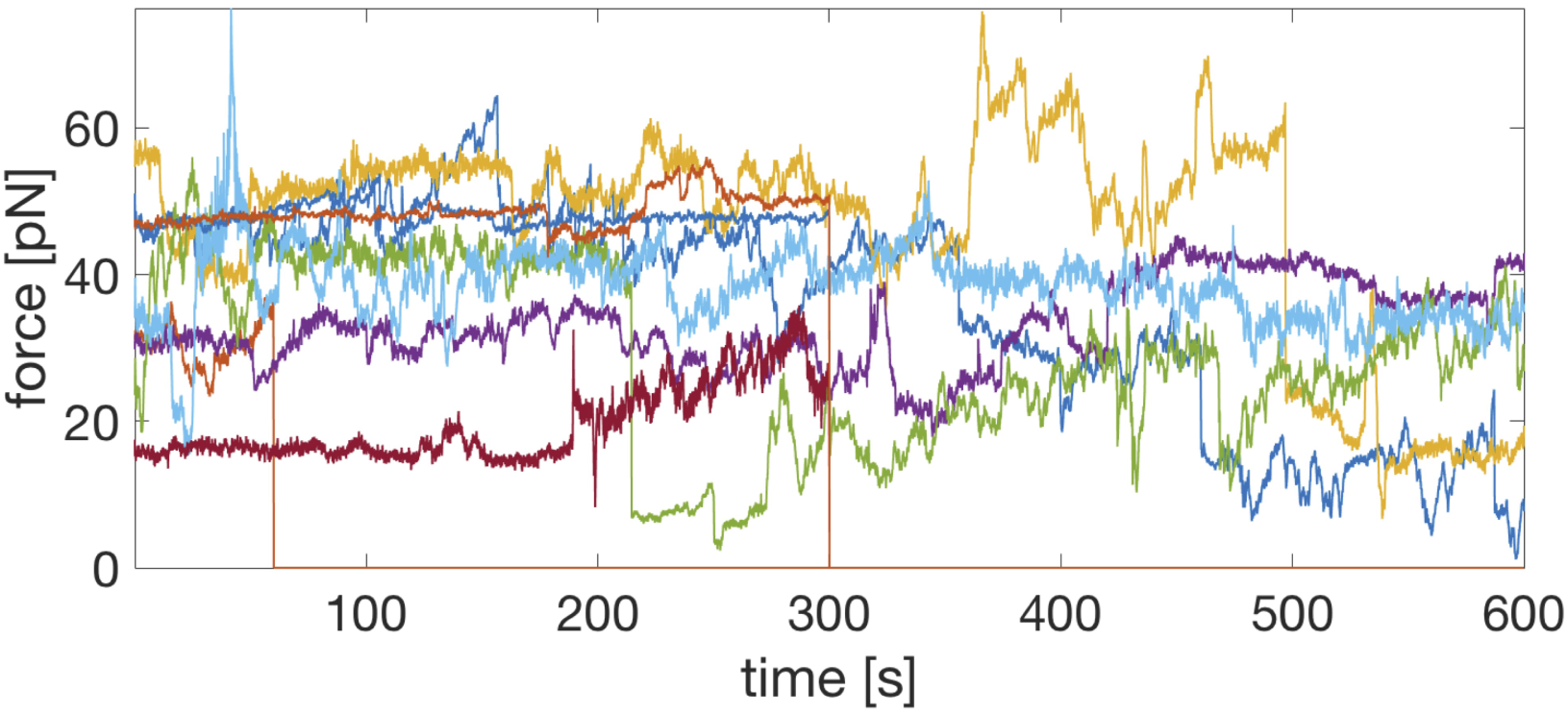
Tethers extracted from MCF7 cells show rich dynamics. 9 different force curves of tethers from MCF7 cells extracted with and held by an optically trapped bead (*d* = 4.95 *μ*m). Acquisition of the forces started after tether extraction.

**Table S5C.** Mean maximum forces for the data in Figure 5F.

| cell type | N | (mean maximum force ± std.) [pN] |
| --- | --- | --- |
| **HEK293** | 60 | $43.8783 \pm 27.1404$ |
| **MCF7** | 11 | $61.0963 \pm 17.8468$ |
| **mESC (2i)** | 42 | $31.3262 \pm 12.7301$ |
| **mESC (S/L)** | 51 | $25.5621 \pm 15.4024$ |

**Table S5D.** *p*-values for the maximum tether force data from Figure 5F. * *p* ≤ 0.05, ** *p* ≤ 0.01, *** *p* ≤ 0.001. The *p*-values for the not mentioned couples show no significant difference between the compared populations.

| cell types | *p*-value |
| --- | --- |
| MCF7, mESC (2i) | $2.68 \cdot 10^{-8}$ *** |
| MCF7, mESC (S/L) | $5.63 \cdot 10^{-9}$ *** |
| HEK293, mESC (2i) | 0.0071 ** |
| HEK293, mESC (S/L) | $7.87 \cdot 10^{-5}$ *** |

**Fig. S6.**
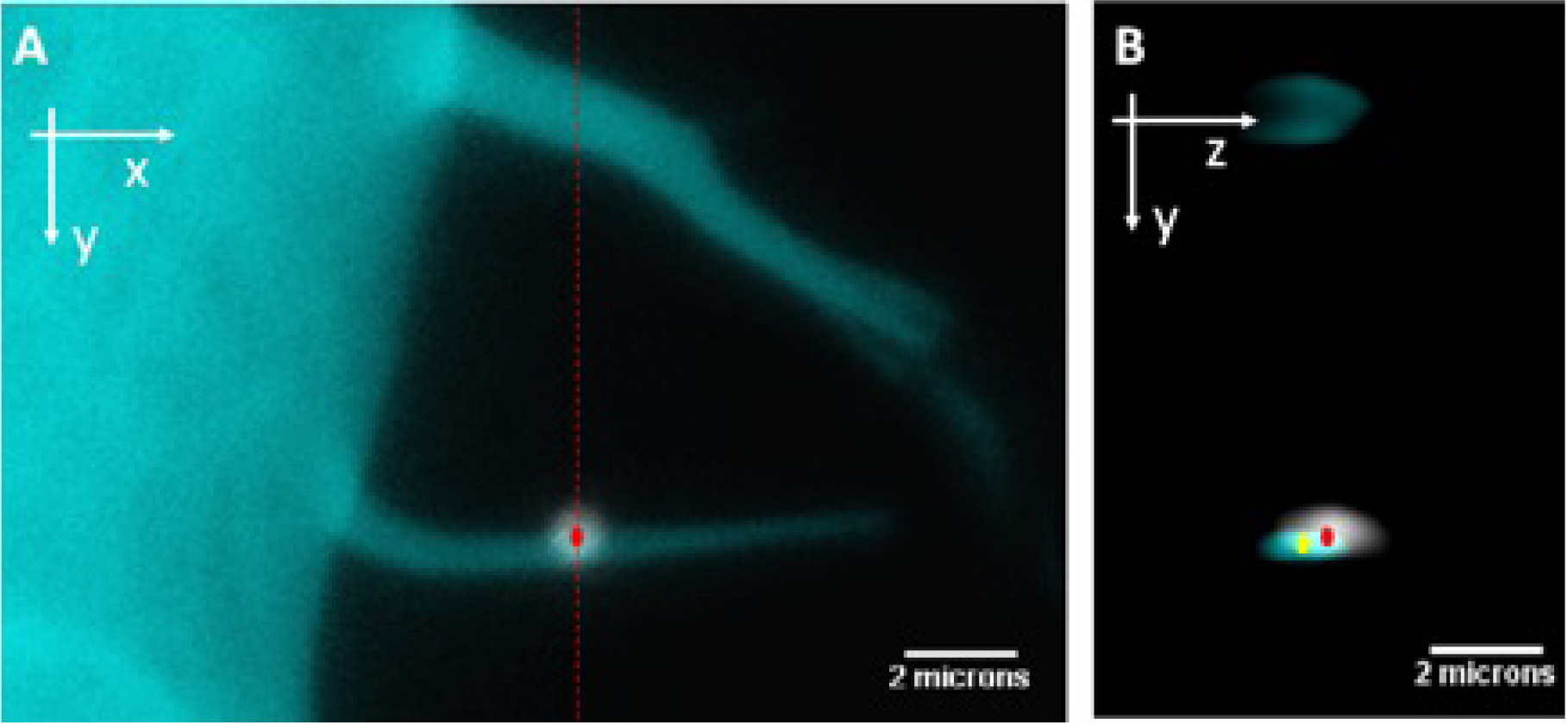
Tracking the bead movement along a filopodium. (A) The *xy*-position of a bead on a filopodium was extracted from the *z*-projection image (red point). (B) The orthogonal view along red dashed line in (A) was used to localize the *yz*-positions of the bead (red point) and center of the filopodium (yellow point).

**Fig. S7.**
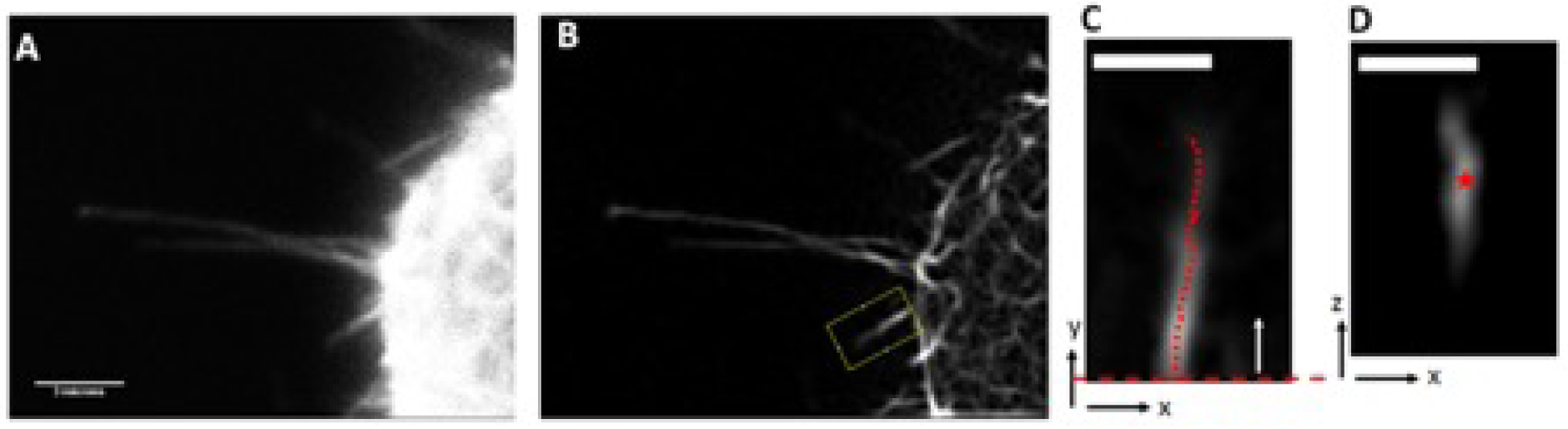
Filopodial tip tracking. (A) *z*-projection of 3-dimensional volume of raw confocal images of a cell with filopodia. (B) The same image from (A) after applying linear Gabor filter to enhance the resolution. (C) The *xy* position of the desired filopodium (yellow box in (B)) was found by scanning the red dashed line along filopodium and applying a Gaussian fit to the intensity profile. (D) The *z* location of the filopodium in each point (red points in (C)) was extracted from the orthogonal view of the image.

**Fig. S8.**
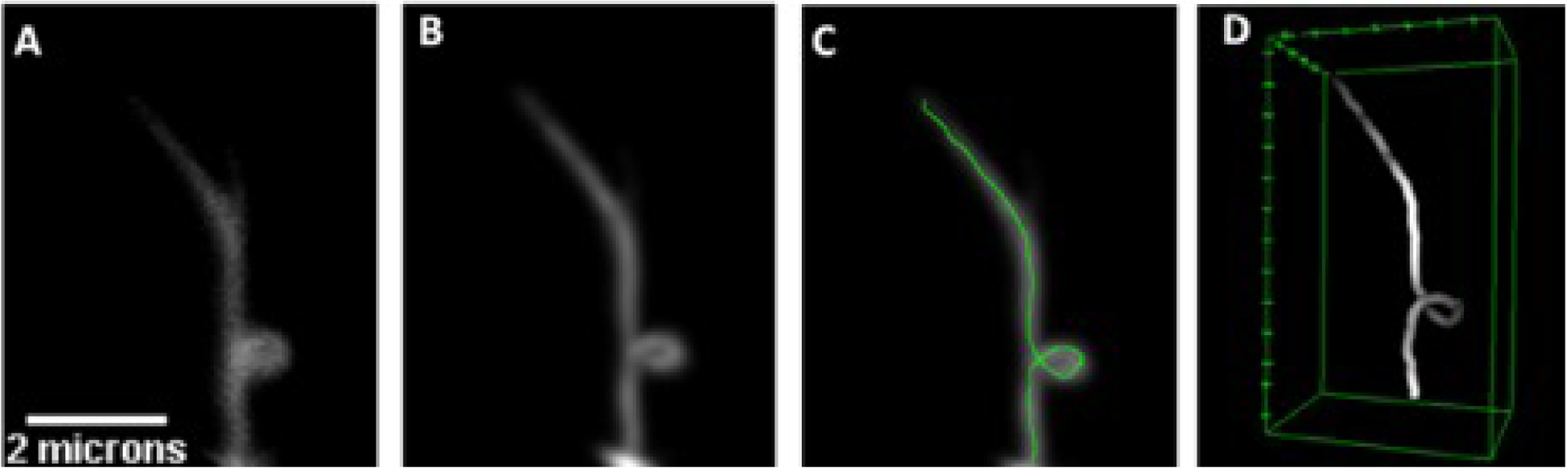
Tracking helical coils. (A) *z*-projection of the 3-dimensional volume containing a coiling filopodium. (B) Gabor and Gaussian blur filtered result of the image from (A). (C and D) Segmentation of the filopodium in 2D (C) and in a 3D (D) view; both obtained using the “Simple Neurite Tracer” plugin in ImageJ.

**Fig. S9.**
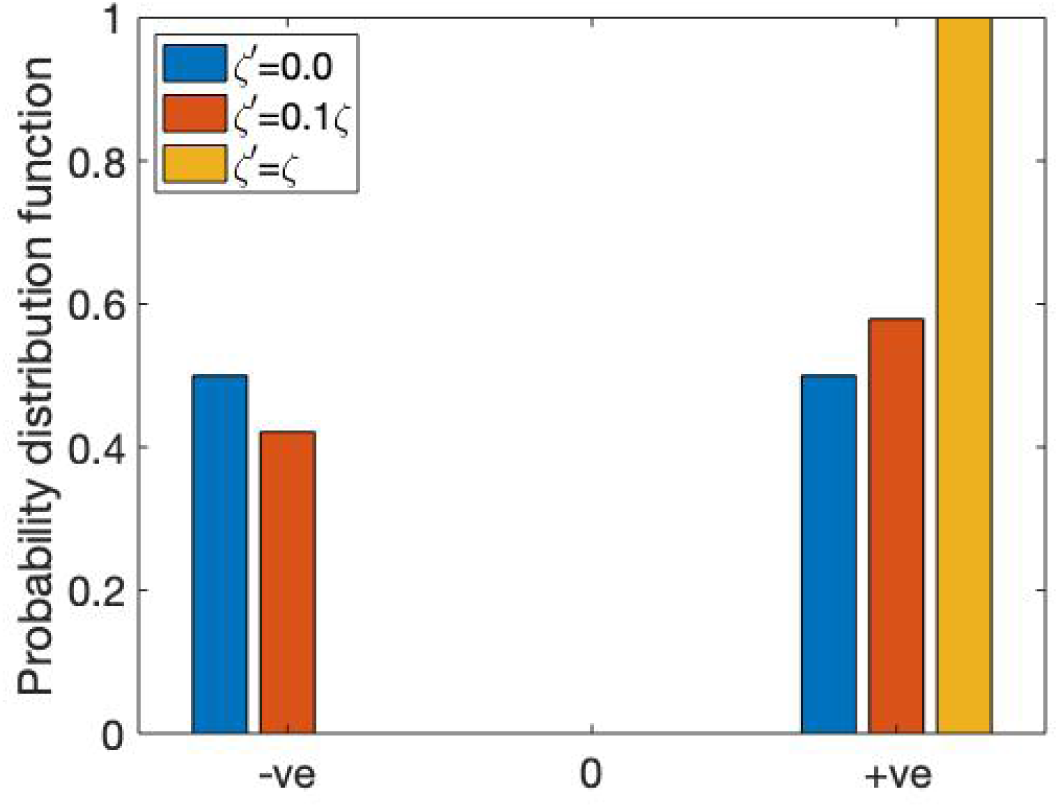
Probability distribution functions for the direction of rotation. Increasing the magnitude of the active torque dipole *ζ*′ increases the biased rotation. Positive (negative) values are indicated by +*ve* (−*ve*).

## 5 Supplementary Movies

- **SI movie 1a HEK293T filopodia sweeping.avi** Sweeping filopodia from the HEK293T cell (cyan, EGFP Lifeact-7) from Figure 1A,B over a time span of 400 s. Scale bar is 5 *μ*m.
- **SI movie 1b KPC 4mgml collagen I bending arr.avi** Bending filopodia from the KPC cell (cyan, EGFP Lifeact-7) from Figure 1C,D and Figure S4a in 4 mg/ml collagen I (gray, reflection) over a time span of 475 s. Red arrows highlight bending regions. Scale bar is 5 *μ*m.
- **SI movie 1c MCF7 tether force.avi** Bending tether pulled from the MCF7 cell (cyan, EGFP Lifeact-7) from Figure 1E (rotated) using a VN coated 4.95 *μ*m bead (gray, reflection). Movie corresponds to the force curve in Figure 1G. Scale bar is 5 *μ*m.
- **SI movie 2a bead rotation MCF7-p95ErbB2.avi** 3D reconstruction of confocal *xyzt* data for the data from Figures 1C-G. VN coated 0.989 *μ*m bead (cyan, flash red) rotating counterclockwise (as seen from tip towards cell body) around a pulled filopodium from a not activated MCF7-p95ErbB2 cell (red, EGFP Lifeact-7). The filopodium is held by a 4.95 *μ*m VN coated bead (gray, reflection) which is glued to the sample surface. Frame interval is 8 sec, total time is 384 s.
- **SI movie 3a mESC filopodia 2i.avi** Filopodia on a mESC (cyan, EGFP Lifeact-7) grown in 2i medium.
- **SI movie 3b hepa cell SiR actin filopodia rotation arr.avi** Rotating filopodia on Hepa 1-6 cell (red, SiR-actin) over 71.5 min, highlighted via a white arrow. Scale bar is 10 *μ*m.
- **SI movie 3c hepa cell SiR actin filopodia rotation arr.avi** Rotating filopodia on Hepa 1-6 cell (red, SiR-actin) over 58 min, highlighted via a white arrow. Scale bar is 10 *μ*m.
- **SI movie 4a KPC 4mgml collagen I bending arr.avi** Bending filopodia from KPC cell (cyan, EGFP Lifeact-7) in 4 mg/ml collagen I (gray, reflection) over a time span of 399 s. Red arrows highlight bending regions. Scale bar is 10 *μ*m.
- **SI movie 4b MCF7 coil.avi** Coiling filopodium from the MCF7 cell (cyan, EGFP Lifeact-7) from Figures S4b, 7C,D over 160 s. Scale bar is 5 *μ*m.
- **SI movie 5a MCF7 extraction of bending tether.mov** Tether pulling from a MCF7 cell (cyan, EGFP Lifeact-7) using a VN coated 4.95 *μ*m bead (grey, reflection). After extraction, the image zoomed in and the focus is adjusted such that the tether is aligned with the imaging plane. Total time is 450 s. Scale bar is 5 *μ*m.
- **SI movie 5b mESC tether.avi** Bending tether pulled from a mESC (cyan, EGFP Lifeact-7) using a VN coated 4.95 *μ*m bead (grey, reflection).

